# Female iPSC X-chromosome inactivation (XCI) erosion and its transcriptomic effects during CRISPR gene editing and neural differentiation

**DOI:** 10.64898/2026.02.27.708613

**Authors:** Christina Thapa, Emily K. Oh, David Sirkin, Jennifer Lahey, Sol Díaz de León Guerrerro, Ada McCarroll, Prarthana Gowda, Hanwen Zhang, Alexandra Barishman, Lilia Peyton, Siwei Zhang, Rebecca M. Pollak, Ronald P. Hart, Carlos N. Pato, Anat Kreimer, Jennifer G. Mulle, Alan R. Sanders, Zhiping Pang, Jubao Duan

**Affiliations:** Center for Psychiatric Genetics, Endeavor Health, Evanston, IL, USA; Department of Psychiatry and Behavioral Neuroscience, The University of Chicago, Chicago, IL, USA; Department of Neuroscience and Cell Biology, Rutgers Robert Wood Johnson Medical School, New Brunswick, NJ, USA; Center for NeuroMetabolism, Child Health Institute of New Jersey, Rutgers Robert Wood Johnson Medical School, New Brunswick, NJ, USA; Center for Advanced Biotechnology and Medicine, Rutgers Robert Wood Johnson Medical School, Piscataway, NJ, USA; Department of Psychiatry, Rutgers Robert Wood Johnson Medical School, New Brunswick, NJ, USA; Department of Cell Biology and Neuroscience, Rutgers University, Piscataway, NJ, USA; Department of Biochemistry and Molecular Biology, Rutgers, The State University of New Jersey, Piscataway, NJ, USA

**Keywords:** Human pluripotent stem cells (hiPSC), *X-chromosome inactive specific transcript* (XIST), X-chromosome inactivation (XCI), neural differentiation, CRSIPR gene editing, neurodevelopment, XCI erosion, autosomal, allele-specific expression (ASE)

## Abstract

Human induced pluripotent stem cells (hiPSC) and iPSC-differentiated neural cells, in combination with CRISPR editing, are commonly used for studying neurodevelopmental and other brain disorders. Female iPSCs undergo random X-chromosome inactivation (XCI) via epigenetic silencing by noncoding *X inactive specific transcript* (*XIST*). It is known that female iPSCs may lose *XIST* expression, leading to XCI erosion that affects both X-linked and autosomal gene expression. However, the effects of CRSIPR editing and neural differentiation on XCI erosion in iPSC-derived neurons and how this may confound a real-world transcriptomic analysis of differentially expressed genes (DEGs) are poorly understood. Here, leveraging bulk RNA-seq of hundreds of CRISPR-edited female iPSC lines from four donor lines for 66 genes and single-cell RNA-seq of iPSC-derived neurons of a subset of 42 edited genes, we investigated the effects of XCI erosion during CRISPR editing and in iPSC-derived neurons. We found that XCI erosion was variable in CRISPR-edited female iPSCs and largely preserved in iPSC-derived neurons. Like in iPSCs, *XIST* in neurons predominately influenced the expression of X-linked genes; however, its effect on autosomal genes was more pronounced in single neurons. Mechanistically, *XIST* epigenetically causes allelic imbalance of both X-linked and autosomal genes, with the former showing stronger allele-specific expression (ASE) bias. Notably, *XIST-*induced ASE bias exhibited a conserved positional pattern at loci affecting neurodevelopmental genes across different female lines and cell types. Finally, we demonstrated a confounding effect of XCI erosion on DEG analyses in iPSC-derived neurons. These results have significant implications in hiPSC modeling of neurodevelopmental and other brain disorders.

## INTRODUCTION

Female cells (X/X) are well known to undergo random X-chromosome inactivation (XCI), resulting in dosage compensation between the sexes. The key regulator of XCI is a long non-coding RNA called *X-chromosome inactive specific transcript* (*XIST*). It coats one of the X-chromosomes and induces transcriptional silencing by recruitment of various RNA-binding proteins and chromatin modifiers^1–3^. This permanent epigenetic change occurs during the blastocyst stage at approximately 4-6 days after fertilization in humans and is maintained in all somatic cells. Although XCI in females is permanent *in vivo*, *XIST* expression in cultured human embryonic stem cells (hESCs) and induced pluripotent stem cells (hiPSCs) has been reported to be highly dynamic. Studies have shown that hiPSCs and hESCs often gradually lose *XIST* expression after a certain number of cell passages or under specific conditions, leading to XCI erosion^4–11^. Recent studies in naive human pluripotent cells and in hiPSCs have shown that *XIST* not only represses X-linked genes as expected, but also spreads a repressive effect to some autosomal regions^12,13^. XCI erosion in females thus has profound effects on iPSC reprogramming, cell differentiation, and disease modeling in the context of widely used CRISPR/Cas9 gene editing in iPSCs.

While XCI erosion is well known in iPSCs, the effects of genome editing and cellular differentiation on XCI erosion dynamics remain poorly understood. In fact, CRISPR-mediated gene editing of iPSCs is a lengthy process spanning multiple cellular passages, and differentiation of iPSC cells into different cell types may further alter *XIST* expression, potentially influencing the regulation of X-linked and autosomal genes. Emerging evidence suggests that gene reactivation associated with XCI erosion in hiPSCs can persist through trilineage and cardiomyocytes differentiation^14^. Whether similar stabilization occurs during neuronal differentiation, particularly in the context of genome editing, remains unclear. Given that hiPSC-differentiated neural and glial models are increasingly used for studying neurodevelopmental and neurodegenerative disorders^15–21^, it is imperative to understand XCI erosion dynamics in hiPSCs during CRISPR gene editing and in neuron/glia differentiation and to determine which X-linked and/or autosomal genes expression are affected, as both the occurrence of erosion and the degree to which genes are reactivated or silenced can vary across cells and genes.

While global effects of XCI erosion on X-linked and autosomal genes are commonly ascertained by examining average expression levels of X-linked or autosomal genes^12^, allele-specific expression (ASE; i.e., allelic imbalance of gene expression) analysis^22^ has been used to identify specific genes that may be influenced by *XIST*^13,14^. ASE analysis allows for the detection of imbalanced expression between the two alleles at heterozygous single nucleotide polymorphism (SNP) sites, providing insights into *cis*-regulatory mechanisms. Recent studies have shown that a modest decrease in *XIST* expression increases biallelic expression (i.e., less ASE bias) of X-linked genes^13,23^. However, the effect of XCI erosion on ASE bias of X-linked and autosomal genes in iPSC-derived neurons has not been systematically investigated. Moreover, whether the genomic loci showing *XIST*-induced ASE biases are conserved across different genetic backgrounds remains unknown.

Recently, as part of the Scalable and Systematic Neurobiology of Psychiatric and Neurodevelopmental Disorder Risk Genes (SSPsyGene) consortium, we have been utilizing CRISPR-based cytosine base editors (CBEs) to establish loss-of-function (LoF) mutagenesis by introducing a premature stop codon (changing “C” to “T” in DNAs) for a large number of neurodevelopmental and psychiatric disorder (NPD)-associated genes in iPSCs from multiple donor lines^19,24,25^. The edited isogenic lines and the non-targeting control (NTC) lines (unedited) are differentiated into both excitatory and inhibitory neurons^19^. These CRISPR-edited iPSC lines for many LoF genes and the respective iPSC-derived neurons provide a unique opportunity for assessing XCI erosion dynamics during CRISPR editing of iPSCs and neuronal differentiation. Here, we investigated the effects of XCI erosion in CRISPR-edited female iPSCs for 66 genes across six donor lines (including 4 female lines) and in iPSC-derived excitatory and inhibitory neurons^19^ for 42 genes along with their matched unedited control lines. We assayed the transcriptomic profiles of iPSC lines and their derived neurons co-cultured with mouse astrocytes by using bulk RNA-seq and single-cell RNA-seq (scRNA-seq)^26^ (Figure 1A, Supplementary Table 1, see methods), respectively. We found that XCI erosion was widespread in CRISPR-edited female iPSC lines and the erosion status of *XIST* expression was largely maintained in iPSC-derived neurons. Like in iPSCs, *XIST* predominantly influenced the expression of X-linked genes in iPSC-derived neurons; however, the repressive transcriptional effect of *XIST* on autosomal genes was more pronounced in single neurons and in the allelic imbalance of expression. We also showed that *XIST-*induced ASE bias on various loci was conserved across different female donor lines and between iPSCs and neurons, likely affecting certain neurodevelopmental genes. Finally, we demonstrated the non-negligible confounding effects of XCI erosion on a real-world differential gene expression analysis in iPSC-derived neurons.

**Figure 1:**
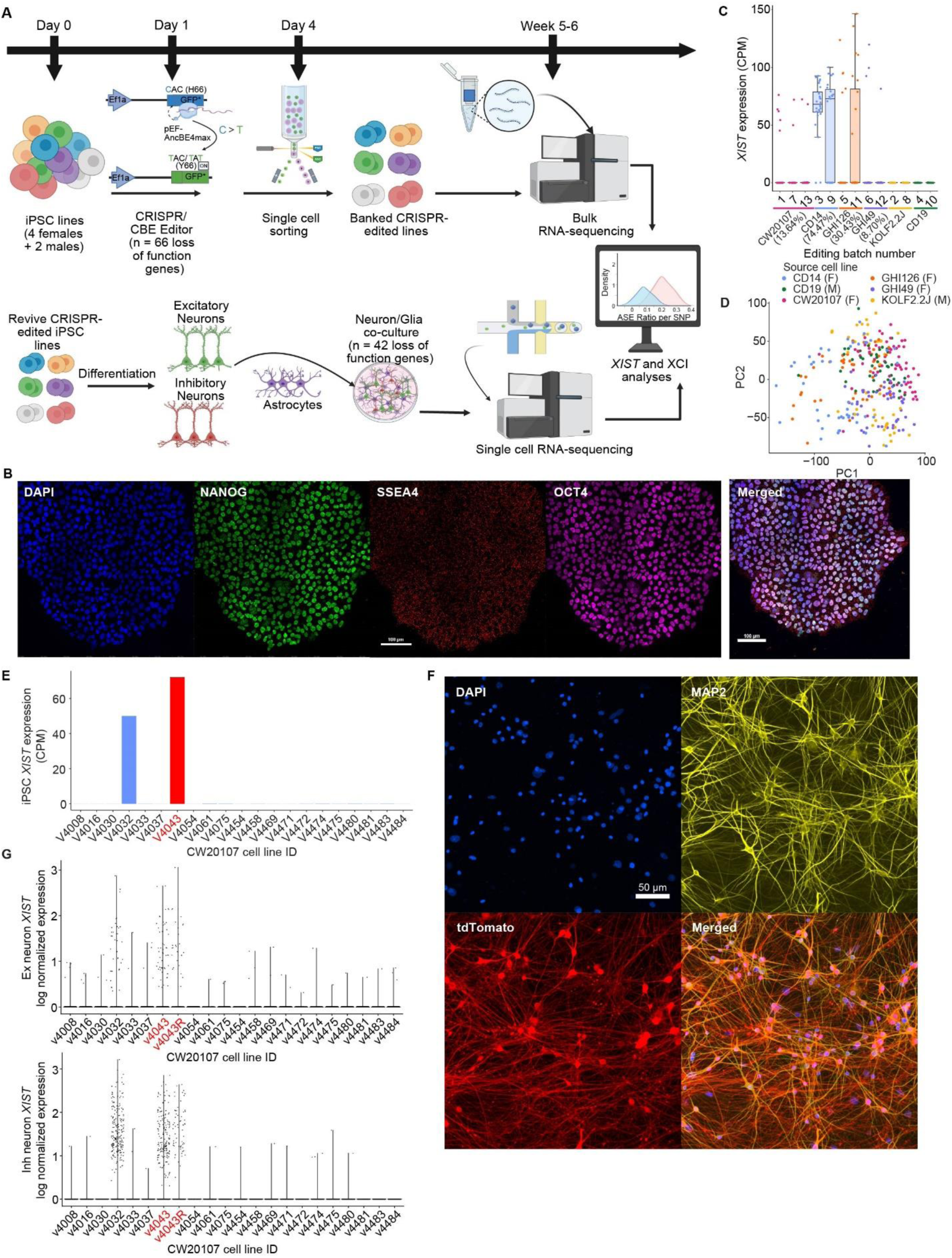
RNA-seq reveals that XCI erosion in female iPSC lines is preserved during iN differentiation. (A) Schematic of experimental design. Six donor lines were used for CRISPR-based DNA editing of 66 target genes to introduce a premature stop codon (iSTOP) in batches (∼23 genes and one donor line per batch). The isogenic iPSC lines for 42 edited genes (both edited and unedited) were differentiated into excitatory and inhibitory neurons and co-cultured with mouse astrocytes. The iPSC and iPSC-derived neuron/glia co-cultures were subject to bulk RNA-seq and scRNA-seq, respectively. (B) Representative immunofluorescence staining of iPSC mutant lines for pluripotent stem cell markers (NANOG, SSEA4 and OCT-4). DAPI, nuclei. Scale bar: 100μm. (C) Box plot showing expression of *XIST* (counts per million reads, CPM) for different editing batches. Each point represents an RNA-seq sample of each line for each batch. Dot color indicates different donor line. The percentage of *XIST*^+^ samples per donor line is indicated in parenthesis. (D) PCA plot of RNA samples of different isogenic iPSC. Color scheme of each dot is the same as in (C). (E) *XIST* expression in different isogenic iPSC lines in CW20107 cell lines of the same batch. Non-targeting control (NTC; line V4043) is shown in red. (F) Immunofluorescence staining of the iPSC-differentiated neurons of day 35. MAP2 staining is for neural dendrites. tdTomato staining is for excitatory neurons (infected by AAV-tdTomato before co-cultured with inhibitory neurons). Scale bar: 50μm. (G) Violin Plot showing *XIST* expression in excitatory (Ex) and Inhibitory (Inh) neurons of different isogenic lines for one example batch using CW20217 as a donor iPSC line for iSTOP base editing. Non-targeting control (NTC; line V4043 and its repeat sample V4043R) is shown in red.

## RESULTS

### XCI erosion is widespread in CRISPR-edited female iPSC lines and largely preserved during neuron differentiation

We first examined the *XIST* expression changes in female iPSC lines after CRISPR DNA base editing. We analyzed bulk RNA-seq data of 297 iPSC lines derived from 6 donor lines (4 female and 2 male), including both the edited and unedited control lines. Immunostaining of pluripotency markers (NANOG, SSEA4, and OCT4) of the CRISPR-edited isogenic lines confirmed their pluripotency following gene editing (Figure 1B). The edited iPSCs from different donor lines exhibited a donor-line-specific pattern of *XIST* expression, with more CD14 isogenic lines showing *XIST* expression compared to the other female lines (Figure 1C). Principal component analysis (PCA) showed no separation of samples based on source cell line, indicating that they remain transcriptome-wide similar to each other despite variable XCI erosion across all female lines (Figure 1D). We observed varying levels of XCI erosion across CRISPR-edited isogenic lines of the same donors (Figure 1E, Supplementary Figure 1A). These iPSC lines had cell passage numbers ranging from 19 to 26; however, there was no significant difference in *XIST* expression between different passages, even for the same donor line (e.g., CW20107) (Supplementary Figure 2A). From comparing NTCs (i.e., iPSCs which went through the same editing process as other LoF mutant lines but with no base editing reagents added) of the same donor line, we observed that editing did not significantly affect *XIST* expression (Supplementary Figure 2B). Thus, the variable XCI erosion across the CRISPR-edited lines from the same female donor line can likely be attributed to heterogeneity of *XIST* expression in the source iPSC population.

We next conducted *XIST* erosion analysis in excitatory and inhibitory neurons differentiated from a subset of these iPSC lines (n=234 lines from 6 donor lines; two were males for comparison purpose) (Figure 1F). Successful differentiation of neurons was confirmed by immunostaining for MAP2, a pan-neuronal marker (Figure 1F). scRNA-seq of these neurons confirmed that in most cases, expression of *XIST* in female iPSCs was preserved in neurons differentiated from these cells; conversely, when *XIST* was not expressed in iPSCs, the derived neurons also lacked *XIST* expression (Figure 1E, G; Supplementary Figure 1). However, in some instances, we did observe that neurons differentiated from the iPSC lines without *XIST* expression (e.g., V4502 in batch 3) regained *XIST* expression (Supplementary Figure 1), suggesting a stochastic nature of *XIST* expression regulation during neuron differentiation from iPSCs.

To examine whether *XIST* expression levels may influence iPSC differentiation into neurons, we calculated pluripotency scores of these iPSC lines from their RNA-seq data and examined their neuronal differentiation capability. Although we found that higher *XIST* expression appeared to be significantly associated with lower pluripotency scores, the differences of pluripotency scores were very subtle (from 0.95 to 0.94) (Supplementary Figure 2C). Likewise, we observed no correlation between *XIST* expression and arbitrarily defined “difficulty” scores of neuron differentiation (Supplementary Figure 2D).

### XCI erosion elevates X-linked and mitochondrial gene expression in CRISPR-edited iPSCs

In human pluripotent stem cells, *XIST* has been shown to repress the expression of X-linked genes and induce gene expression dampening across specific autosomal regions^12^. Here, to test for the effect of XCI erosion on X-linked genes and a broader transcriptional effect in iPSC lines during CRISPR editing, we first compared the mean expression of X-linked and autosomal genes among female iPSC lines (with or without *XIST* expression, *XIST*^+^ or *XIST*^-^) compared to that of male iPSC lines. As expected, we found that the *XIST*^-^ female iPSC lines had higher expression of X-linked genes compared to *XIST*^+^ female lines and male lines, confirming the effect of XCI erosion for the *XIST*^-^ group (Figure 2A, Supplementary Figure 2E, 2G). Interestingly, we also observed a subtle increase of X-linked gene expression in *XIST*^+^ female lines vs. male lines, suggesting a possibly incomplete XCI in those *XIST*^+^ female lines (Figure 2A). However, for autosomal genes, in contrast to prior study of hiPSCs^12^, we found lower expression of autosomal genes in *XIST*^-^ female iPSC lines compared to the other groups, though, it was a subtle difference of expression (Figure 2B, Supplementary Figure 2F, H).

**Figure 2:**
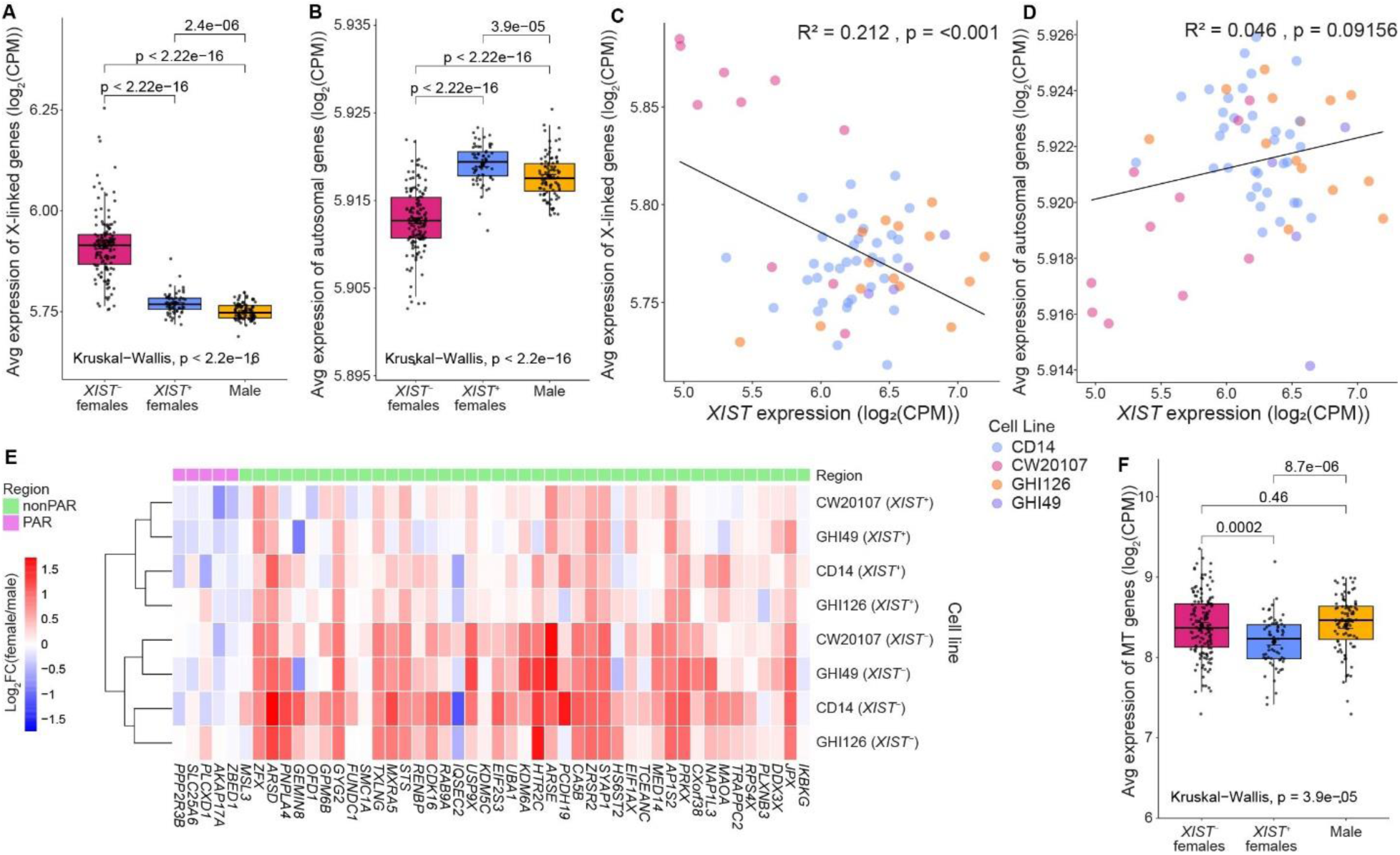
Transcriptomic effects of *XIST* erosion in iPSCs. (A-B) Box-and-whisker plots comparing the expression of (A) X-linked genes and (B) autosomal genes between male, *XIST*^+^ females, and *XIST*^-^ females. The box represents the interquartile range (IQR) with the median shown as the central line. Whiskers extend to the furthest data points within 1.5 × IQR from the quartiles. Each dot represents an isogenic cell line. (C-D) Scatter plots showing the correlation of *XIST* expression in a cell line to average expression of (C) X-linked genes and (D) autosomal genes. CPM, counts per million reads. (E) Heatmap showing the expression levels (normalized to male) of the reported XCI-escape genes (N = 48) in female isogenic iPSC lines with different XCI status. PAR, pseudoautosomal regions of X-chromosome. (F) Box-and-whisker plot comparing the expression of mitochondrial (MT) genes (encoded by MT genome) between male, *XIST*^+^ females, and *XIST*^-^ females. Box represents interquartile range and whisker represents 1.5 × IQR from the quartiles. Each dot represents an isogenic cell line.

To further corroborate the above-described effects of *XIST* expression on X-linked and autosomal genes, we performed linear regression analysis on *XIST*^+^ isogenic lines using *XIST* expression as a predictor and average gene expression as the outcome. We found a modest negative correlation between *XIST* expression and X-linked gene expression, with *XIST* expression explaining ∼20% of the variance in X-linked gene expression (R^2^ = 0.21, *p* < 0.01) (Figure 2C). In contrast, no significant association was found between *XIST* and autosomal gene expression (R^2^ = 0.04, *p* > 0.09) (Figure 2D). Together, our data shows that the regulatory effect of *XIST* is largely specific to X-chromosome genes in CRISPR-edited iPSCs.

Studies have shown that over 15% of X-linked genes escape *XIST*-mediated inactivation and remain transcriptionally activated even on the inactive X-chromosome across human tissues^23,27,28^. We hypothesized that the absence of *XIST* may also lead to increased expression of these XCI-escaping genes in our iPSC lines. To investigate this, we examined the expression pattern of known XCI-escaping genes across different *XIST*^+^ and *XIST*^-^ isogenic lines (Figure 2E). Sex chromosomes include PARs (pseudoautosomal regions), which are terminal regions of the chromosomes that are autosome-like in that these homologous sequences at the tips of the X-and Y- chromosomes are recombined during meiosis^29^. Consistent with a previous study, we found that genes in the PARs showed higher expression in males compared to females, suggesting that the presence or absence of XCI is unable to overcome male-biased expression^23^. In non-PARs, we see that most of the known XCI-escaping genes indeed show robust expression even in *XIST*^+^ iPSC lines, which is consistent with what is reported across tissues in humans. Furthermore, for many of these XCI-escaping genes, XCI erosion in *XIST*^-^ samples further increased their expression (Figure 2E). Together, these results suggest that the effects of XCI erosion are primarily detectable in non-PARs, with limited impact on sex-biased expression in the PARs, further supporting the predominant effect of XCI erosion on the expression of X-linked genes rather than autosomal genes in our CRISPR-edited iPSCs.

Emerging evidence also suggests that *XIST* may lead to mitochondrial (MT) dysfunctions^30,31^. We thus examined whether the loss of *XIST* is associated with transcriptional changes in MT gene expression (n = 33 MT genes). We observed that MT gene expression was lowest in *XIST*^+^ female lines, and significantly increased in *XIST*^-^ female lines. Notably, MT gene expression levels in *XIST*^-^ females were comparable to those in males (Figure 2F). This pattern is consistent with a recent study reporting a sex specific bias for heart MT gene expression^32^, where protein coding MT genes were expressed in higher levels in males compared to females in mice. These findings suggest that the loss of *XIST* could lead to de-repression of MT genes in females, resulting in expression levels comparable to those observed in males.

### XCI erosion in iPSCs has a stronger effect on allelic imbalance of expression of X-linked genes than autosomal genes

Our observed lack of a significant effect of XCI erosion on average expression level of autosomal genes may be due to the insensitivity of using average gene expression in detecting more subtle effects of XCI erosion. ASE analysis^22^, by directly comparing the RNA-seq counts of the two alleles of an expressed heterozygous SNP within the same sample, provides a sensitive way to identify target genes regulated by *cis*-regulatory genetic variants or epigenetic factors such as *XIST* (Figure 3A). XCI erosion is expected to make an *XIST*-affected gene to shift from ASE towards biallelic expression (Figure 3A). We thus analyzed SNP ASE patterns across both X-linked and autosomal genes in *XIST*^-^ and *XIST*^+^ female lines. To ensure robust estimation of allelic expression, we restricted the analysis to highly expressed heterozygous SNPs (i.e., total read count ≥ 5 times the number of samples per group; see Methods) for X-linked and autosomal chromosomes. For each donor line, we matched the number of *XIST*^+^ and *XIST*^-^ samples, prioritizing keeping the total RNA-seq read depth balanced between groups to ensure comparability. To assess how *XIST* expression influences allelic expression, we categorized the SNPs as biallelic or exhibiting allelic ASE based on the results of a binomial test. We observed substantial overlap in ASE SNPs between *XIST*^+^ and *XIST*^-^ samples, regardless of chromosome type (Figure 3B). This suggests that most SNPs with allelic bias are not uniquely associated with *XIST* status, although the magnitude of allelic imbalance at shared SNPs may differ between *XIST*^+^ and *XIST*^-^ lines. However, when focusing specifically on SNPs classified as biallelic in *XIST*^-^samples, we found that a large proportion of these X-linked SNPs became allelically biased in *XIST*^+^ samples (Figure 3C). This shift toward allelically biased expression in the presence of *XIST* is consistent with re-establishment of XCI, reflecting the fact that XIST⁺ cells maintain a partially inactive X-chromosome that still contributes low-level transcription. In contrast, only a small proportion of autosomal biallelic SNPs in *XIST*^-^ samples transitioned to ASE in *XIST*^+^ samples (Figure 3C), which was consistent with our observed predominant effect of *XIST* expression on X-linked genes (Figure 2).

**Figure 3:**
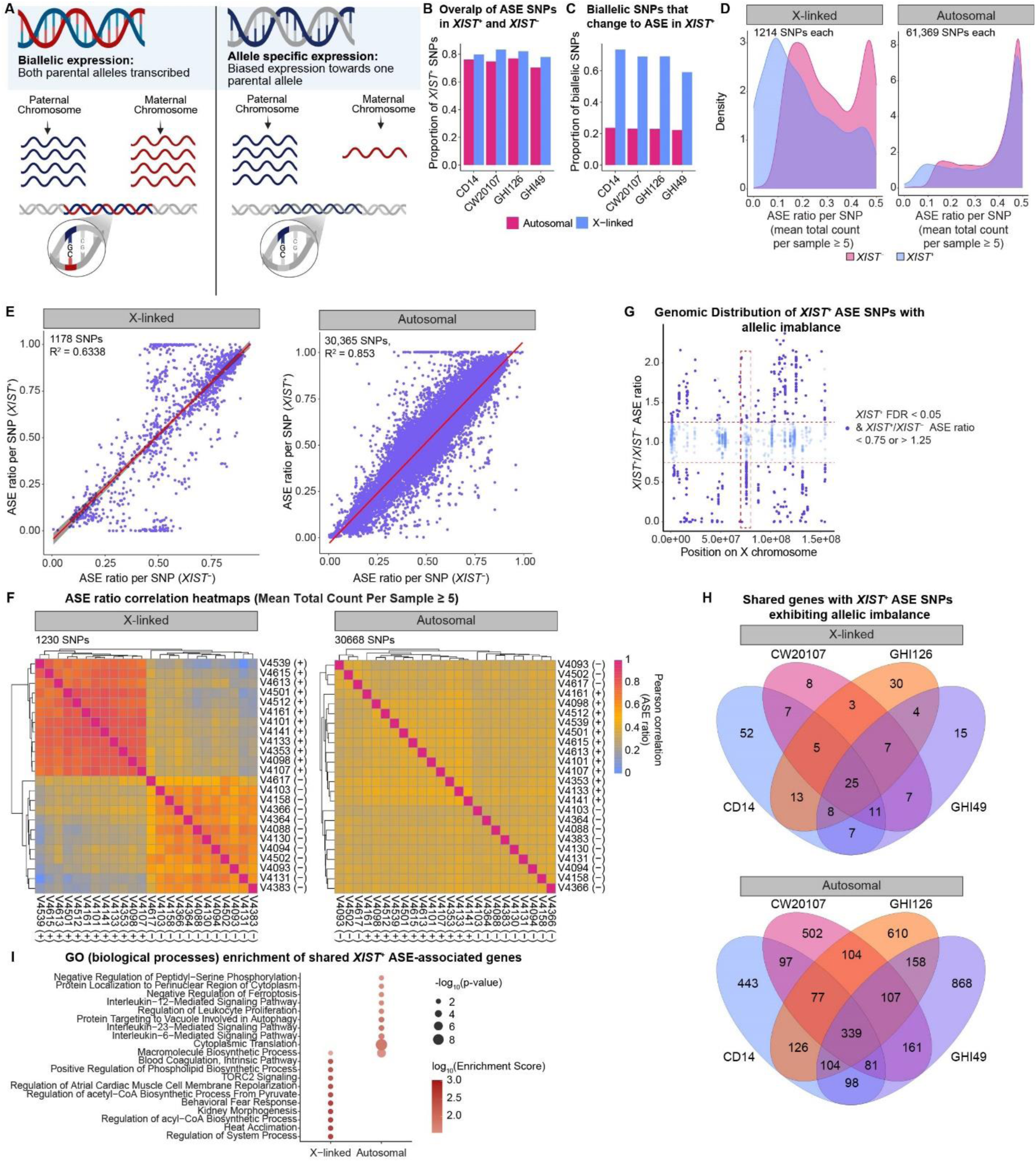
Allelic imbalance of gene expression caused by XCI erosion in female iPSC lines. (A) Schematic representation of biallelic (paternal and maternal alleles) and allele-specific expression (ASE) at a SNP site. (B) Bar plot showing overlap of ASE SNPs in XIST+ and XIST-samples for different female iPSC donor lines. (C) Bar plot showing proportion of biallelic SNPs that change to ASE in the presence of XIST expression in different cell lines. (D) Density plot showing the distribution of ASE ratio for X-linked and autosomal SNPs across XIST- and XIST+ in iPSC line CD14. (E) Scatter plot to compare the degree of allelic imbalance between XIST+ and XIST- samples in iPSC line CD14. Each dot represents one SNP. ASE ratio, the proportion of reference allele counts of all sequencing reads at a SNP site. (F) Pearson correlation heatmap comparing ASE ratios (ref Count / total Count) for each SNP across different XCI statuses of isogenic cell lines derived from donor line CD14. (G) Positional scatter plot showing the distribution of XIST+/XIST- ASE ratio for SNPs across the X-chromosome of iPSC line CD14. Horizontal red dashed line highlights SNPs with XIST+/XIST- ASE ratio < 0.75 or > 1.25 whereas vertical red dashed box highlights the 70-80Mb hot spot of allelic imbalance. (H) Venn diagram showing overlap of genes containing SNPs with ASE ratio (XIST+/XIST-) < 0.75 or > 1.25 across cell lines. (I) Gene ontology (GO) enrichment (biological processes) for genes with XIST+ ASE SNPs overlapping between all cell lines. Top 10 enriched GO terms are listed for overlapping X-linked genes (n = 25) and autosomal genes (n = 339) from (H).

To more quantitatively ascertain ASE differences of X-linked and autosomal genes between *XIST*^-^and *XIST*^+^ groups, we first examined the distribution of the ASE ratios (sum of reference allele count / total allele count) within each donor line (Figure 3D, Supplementary Figure 3A). We found that for the X-chromosome, *XIST*^+^ groups revealed a higher density of monoallelic SNPs (ASE ratios closer to 0) compared to the *XIST*^-^ groups, consistent with XCI. Conversely, *XIST*^-^ groups displayed a shift towards biallelic SNPs (minor allelic ratios closer to 0.5), reflecting XCI erosion. Across all four female lines, XCI erosion showed a consistent pattern of having a much weaker but detectable effect on ASE bias of autosomal SNPs compared to SNPs in X-linked genes (Figure 3D, Supplementary Figure 3A). We next examined the ASE ratio of individual SNPs to compare the degree of allelic imbalance between *XIST*^+^ and *XIST*^-^ samples. For X-linked SNPs, although allelic ratios between *XIST*^+^ and *XIST*^-^ samples remained moderately correlated (Pearson’s R² = 0.73), there was increased deviation from the diagonal (y = x) in both directions, indicating heterogeneous allelic shifts consistent with XCI erosion across the X-chromosome in *XIST^+^* samples. In contrast, the ASE ratios of autosomal SNPs exhibited stronger correlation (Pearson’s R² = 0.83) between *XIST*^+^ and *XIST*^-^ groups, reflecting the smaller impact of XCI erosion on autosomal gene expression (Figure 3E, Supplementary Figure 3B).

To evaluate global differences in allelic expression between *XIST*^+^ and *XIST*^-^ samples, we computed the pairwise Pearson correlation of ASE ratios of highly expressed SNPs between different iPSC lines derived from the same donor line. Higher correlations were consistently observed within the same *XIST* status group (*XIST*^+^ or *XIST*^-^) for X-linked SNPs compared to those between *XIST* status groups (Figure 3F, Supplementary Figure 3C), consistent with the above-mentioned effect of *XIST* erosion on ASE bias. The clear separation of the *XIST*^+^ and *XIST*^-^groups also indicated that the iPSC donor line used for CRISPR editing was clonal (i.e., the same paternal or maternal allele showing bias across all derivative iPSC lines), which otherwise would not show strong correlation of SNP ASE bias across different isogenic lines of the same donor line. Compared to X-linked SNPs, correlations of autosomal SNP ASE ratios between *XIST*^+^ and *XIST*^-^ groups did not show clear separation (Figure 3F, Supplementary Figure 3D), again indicating smaller global effect of XCI erosion on autosomal genes. Together, these results illustrate that XCI erosion in *XIST*^-^ CRISPR-edited iPSC lines alter ASE of both X-linked and autosomal genes, but with a much stronger effect on X-linked genes.

Leveraging our SNP ASE data on multiple donor lines, we next investigated whether the SNPs with ASE bias influenced by *XIST* erosion were restricted to specific loci across different genetic backgrounds. We plotted the ratio of ASE values between *XIST*^+^ and *XIST*^-^ samples for each SNP along the X-chromosome. A ratio of 1 indicates equal ASE between the two cell groups, whereas deviations reflect differential allelic expression potentially due to XCI. We found that most loci had an ASE ratio close to 1, indicating balanced allelic expression (Figure 3G, Supplementary Figure 3D). Notably, although loci showing strong ASE bias appeared to be evenly distributed across the X-chromosome, the pattern of ASE bias seems to be similar across different donor lines, suggesting that specific genomic regions of the X-chromosome were influenced by XCI erosion (Figure 3G, Supplementary Figure 3D). To further identify genes in those regions that are consistently affected by *XIST*-associated allelic regulation across different donor lines, we selected SNPs showing at least a 25% difference in allelic expression between XIST^+^ and *XIST*^-^samples (*XIST*^+^/*XIST*^-^ ASE ratio < 0.75 or >1.25) to their corresponding genes for all source cell lines (Supplementary Tables 2, 3). We found that among genes retaining those strong ASE SNPs, 25 (out of 202) X-linked genes were shared across all female cell lines and 339 (out of 3,875) autosomal genes were shared across all female cell lines (Figure 3H). The gene ontology (GO) term enrichment analysis revealed that the *XIST*-influenced X-linked genes were enriched for biological processes related to TORC2 signaling, positive regulation of phospholipid biosynthesis, and acetyl-CoA metabolism (Figure 3I), indicating possible effects on lipid homeostasis, membrane dynamics, and stress response. For *XIST*-influenced autosomal genes, the enriched GO-terms included interleukin-23-mediated signaling, leukocyte proliferation, and cytoplasmic translation (Figure 3I), suggesting possible alterations in immune-related signaling and the machinery involved in protein synthesis.

### XCI erosion in single neurons affects the expression of both X-linked and autosomal genes

There has been a lack of systematic investigation of the functional effects of *XIST* erosion in iPSC-derived neurons, especially at single cell resolution. As part of the SSPsyGene effort^19^, we analyzed scRNA-seq data of 121,317 excitatory (*SLC17A6*^+^) and 69,022 inhibitory (*GAD1*^+^) neurons derived from iPSC lines of 42 CRISPR-edited LoF alleles and unedited (i.e., NTC) controls for 4 female and 2 male donor lines (Figure 4A, B). We found that most female cell lines showed variable neuronal expression of *XIST*, with the majority of neurons lacking detectable *XIST* (*XIST*^+^: 79,788 cells, *XIST*^-^: 110,551 cells) (Figure 4C, Figure 1G, Supplementary Figure 1B).

**Figure 4:**
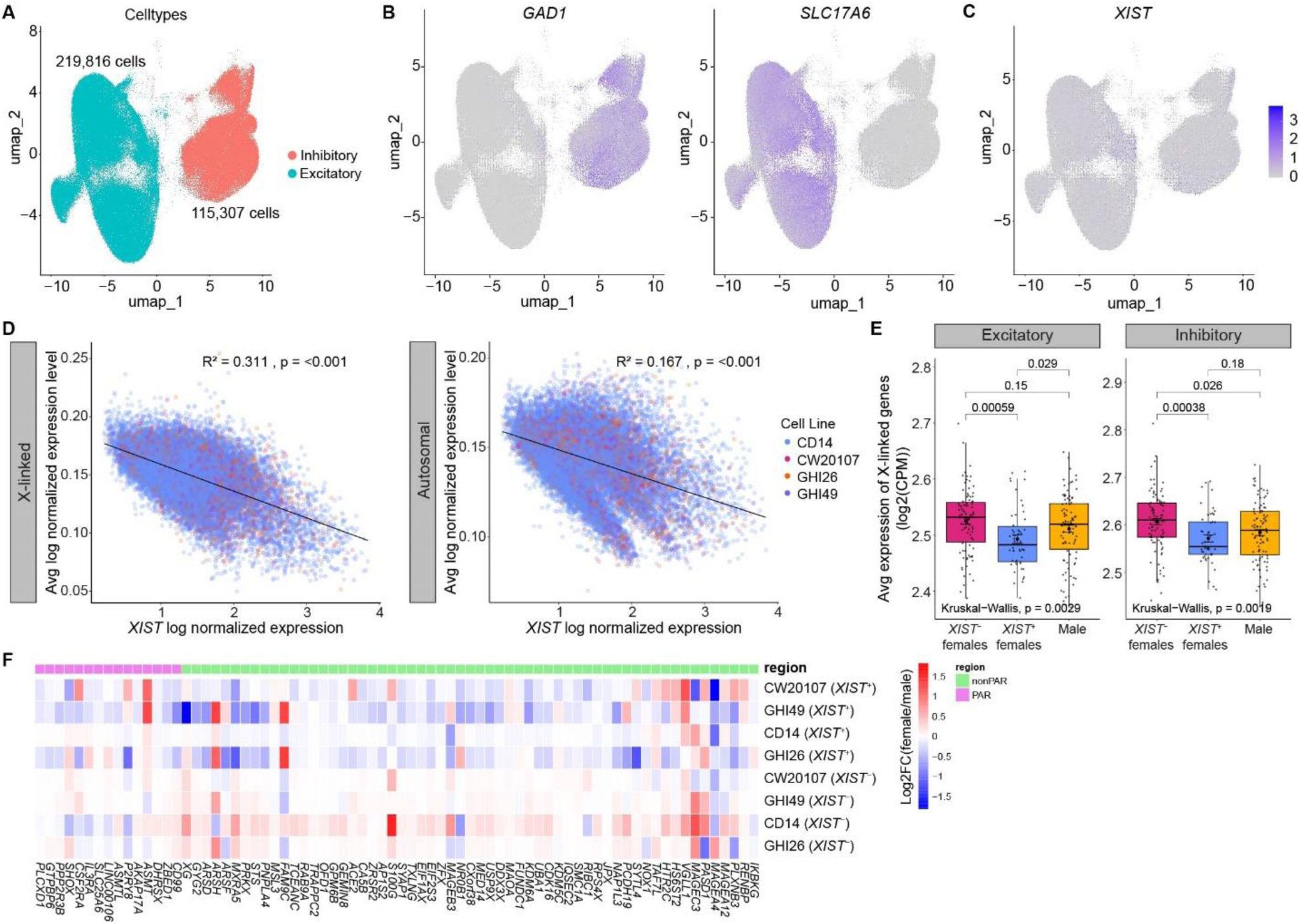
Transcriptomic effects of XCI erosion in female iPSC-derived neurons. (A) UMAP projection of snRNA-seq data of excitatory and inhibitory neurons of 225 isogenic iPSC lines derived from four female donor lines. (B) Feature plots for single cell gene expression of cell type specific markers (SLC17A6 for excitatory and GAD1 for inhibitory). (C) Feature plots for single cell XIST gene expression. (D) Pearson’s correlation of single-cell XIST log-normalized expression and average log-normalized expression of X-linked genes (left) and autosomal genes (right) in excitatory neurons. Each dot is a single cell. Dot color indicates different cell lines. (E) Average expression levels of X-linked in excitatory (left) and inhibitory (right) neurons between XIST- female, XIST+ female, and male lines. Box represents the IQR and whiskers represent 1.5*IQR from the quartiles. Wilcoxson p-value is given. (F) Heatmap showing the expression levels (normalized to male) of the reported XCI-escape genes (N = 74) in excitatory neurons of female isogenic iPSC lines with different XCI status. PAR, pseudoautosomal regions of X-chromosome.

Leveraging our scRNA-seq data, we first examined at single-cell resolution whether XCI erosion affects the expression of X-linked and autosomal genes. By analyzing single cells with detectable *XIST* expression, we found that *XIST* expression showed a moderate effect (R^2^ = 0.31, *p* < 0.01) on X-linked gene expression and a lesser effect (R^2^ = 0.17, *p* < 0.01) on autosomal gene expression in excitatory neurons (Figure 4D). *XIST* expression in inhibitory neurons showed a weaker correlation with X-linked genes (R^2^ = 0.20, *p* < 0.01) but a similar level of correlation with autosomal genes (R^2^ = 0.18, *p* < 0.01) (Supplementary Figure 4A), suggesting a possible neuron subtype-specific effect of XCI erosion in single cells.

We further analyzed the transcriptomic effects of XCI erosion by comparing pseudobulk RNA-expression levels of X-linked and autosomal genes across *XIST*^+^ and *XIST*^-^ female lines as well as male lines. Like in iPSCs, for both excitatory and inhibitory neurons, we observed higher expression of X-linked genes, but not autosomal genes, in *XIST*^-^ female lines (vs. *XIST*^+^ lines) (Figure 4E, Supplementary Figure 4B). As opposed to a significant effect of *XIST* erosion on MT gene expression in iPSCs (Figure 2F), we did not observe an effect of *XIST* expression on MT genes in neurons (Supplementary Figure 4C). Lastly, we examined the expression of known XCI-escaping genes on the X-chromosome in both the inhibitory and excitatory neurons (Figure 4F, Supplementary Figure 4D). We found that unlike in iPSCs (Figure 2F), most known XCI-escaping genes did not “escape” from XCI in *XIST*^+^ female lines, showing relatively low expression (normalized by the expression in male lines) in induced neurons (Figure 4F, Supplementary Figure 4D). However, like in iPSC, *XIST*^-^ neurons did exhibit increased expression of these XCI-escaping genes in the nonPAR, suggesting that XCI erosion still contributes to the de-repression of XCI-escaping genes in neurons (Figure 4F, Supplementary Figure 4D).

Taken together, these results suggest that although XCI erosion does not seem to affect average autosomal gene expression, it is associated with modest transcriptional changes of both X-linked and autosomal genes in single neurons with a stronger effect on X-linked genes.

### XCI erosion in iPSC-derived neurons affects ASE bias of neurodevelopmental genes

Given the observed effects of XCI erosion on ASE bias in iPSC (Figure 3), we also examined whether similar ASE bias can be seen in iPSC-derived excitatory and inhibitory neurons. We first categorized individual cells from each gene’s LoF mutant line into *XIST*^+^ and *XIST*^-^ groups based on detectable levels of *XIST* transcripts. This classification was performed at the single-cell level to ensure accurate representation of *XIST* status across heterogeneous neuronal populations. To minimize the effect of confounding variables, comparable sequencing depths and an equal number of samples were selected for the *XIST*^+^ and *XIST*^-^ groups. Next, to assess the influence of *XIST* expression on allelic bias, we categorized the expressed heterozygous SNPs into biallelic and ASE groups. Like in iPSC, we found that the neuronal ASE SNPs largely overlapped between the *XIST*^+^ and *XIST*- groups across all cell lines and both neuronal subtypes (Figure 5A), and a large proportion of X-linked biallelic SNPs in *XIST*- samples became allelically biased in *XIST*^+^ samples (Figure 5B). In contrast, a smaller proportion of autosomal biallelic SNPs in *XIST*^-^samples transitioned to ASE in *XIST*^+^ samples (Figure 5B). Also like in iPSCs (Figure 3D), density plots of allelic ratios for highly expressed SNPs on the X-chromosome in both types of neurons showed a higher density of monoallelic SNPs (ASE ratios closer to 0) for *XIST*^+^ groups compared *XIST*^-^ groups (Figure 5C, Supplementary Figure 5A). In contrast, for autosomal SNPs, although *XIST*^+^ neurons also exhibited a more pronounced monoallelic expression pattern (centered around allelic ratio 0.1), both the *XIST*^+^ and *XIST*^-^ groups had a bimodal distribution of allelic ratios, with the prominent peak around an allele ratio of 0.5 (i.e., biallelic expression) (Figure 5C, Supplementary Figure 5A). Altogether, these results suggest much stronger transcriptional effects of XCI erosion on X-linked genes compared to autosomal genes in iPSC-derived neurons.

**Figure 5:**
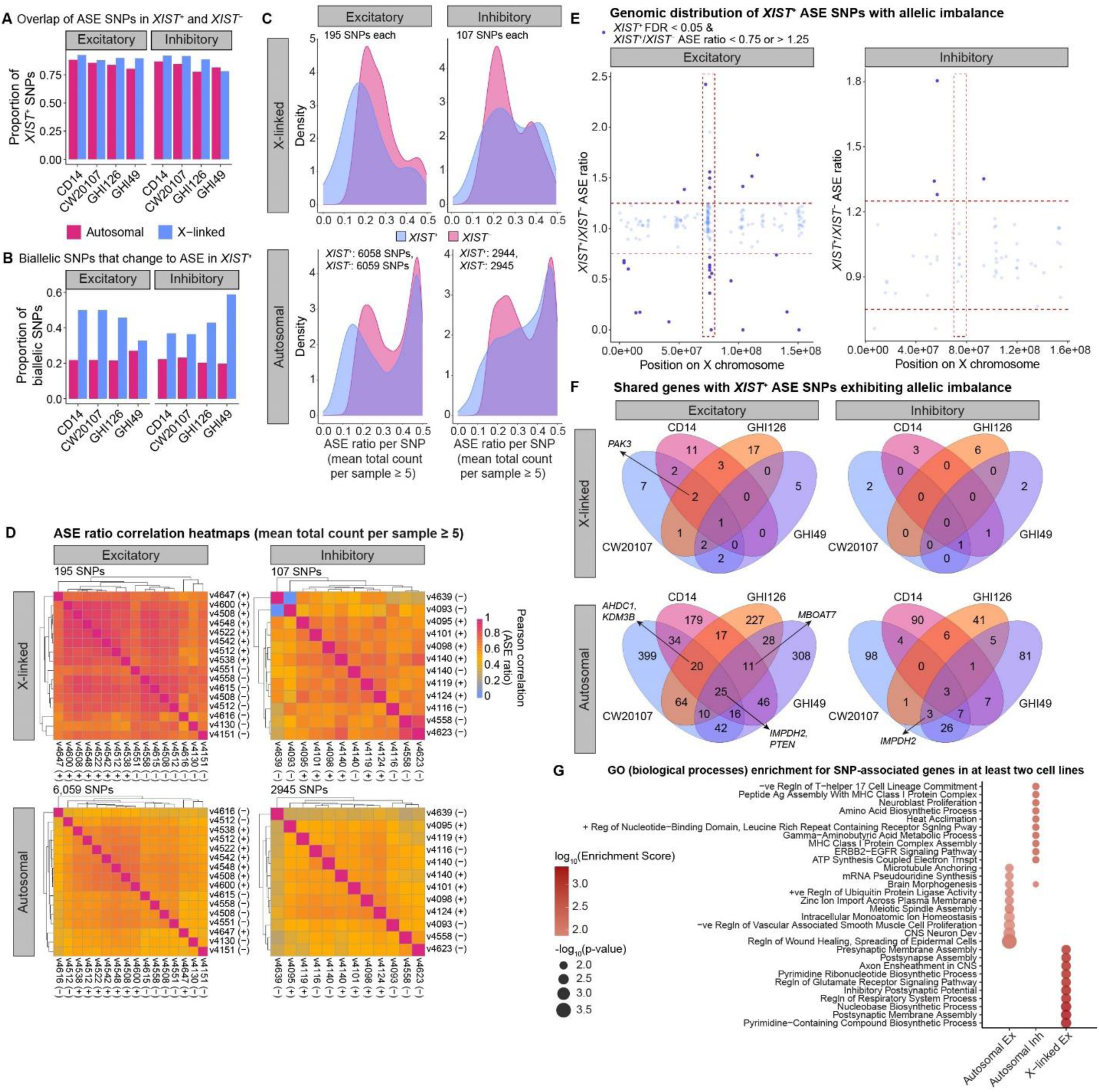
Allelic imbalance of gene expression caused by XCI erosion in female iPSC-derived neurons. (A) Bar plot showing overlap of ASE SNPs in *XIST*^+^ and *XIST*^-^ excitatory (left) and inhibitory (right) neurons for different female iPSC donor lines. (B) Bar plot showing proportion of biallelic SNPs that change to ASE in the presence of *XIST* expression in excitatory (left) and inhibitory (right) neurons for different female iPSC donor lines. (C) Density plot showing the distribution of the ASE ratio for X-linked (top) and autosomal (bottom) SNPs across *XIST*^-^ and *XIST*^+^ in excitatory (left) and inhibitory (right) neurons derived from donor line CD14. (D) Heatmap showing Pearson’s correlation of SNP ASE ratios (ref Count / total Count) of X-linked (top) and autosomal (bottom) genes in excitatory (left) and inhibitory (right) neurons between isogenic lines with different *XIST* status (^+^ or ^-^) derived from the female donor line CD14. (E) Positional scatter plot showing the distribution of *XIST*^+^/*XIST*^-^ ASE ratio for SNPs across X-chromosome of the female donor line CD14. Horizontal red dashed line highlights SNPs with *XIST*^+^/*XIST*^-^ ASE ratio < 0.75 or > 1.25 whereas vertical red dashed box highlights the 70-80Mb hot spot of allelic imbalance. (F) Venn diagram showing overlap of genes containing SNPs with ASE ratio (*XIST*^+^/*XIST*^-^) < 0.75 or > 1.25 for X-linked (top) and autosomal (bottom) genes in excitatory (left) and inhibitory (right) neurons across cell lines. NDD genes overlapping in three or more cell lines are indicated by arrows. (G) Gene ontology (GO) enrichment (biological processes) for genes containing neuronal *XIST*^+^ ASE SNPs overlapping between at least two donor lines. Top 10 enriched GO terms are listed for the overlapping X-linked genes (not available for inhibitory neurons) and autosomal genes from (G).

We also computed the pairwise correlation of ASE ratios to evaluate the global differences in ASE bias between *XIST*^+^ and *XIST*^-^ samples within each cell line and neuronal subtype. In contrast to iPSC data, we did not find evidence of clustering between the same *XIST* status group even for the X-linked genes (Figure 5D, Supplementary Figure 6), suggesting a weaker impact of XCI erosion in neurons than in iPSC.

To investigate positional patterns of XCI-affected SNP ASE bias in neurons across different donors, we plotted the ratio of *XIST*^+^ and *XIST*^-^ ASE values for X-chromosome SNPs. Consistent with iPSC cell lines (Figure 3G), most SNPs were clustered around a ratio of 1, indicating comparable allelic expression between groups (Figure 5E, Supplementary Figure 5B). Despite a smaller number of called SNPs from scRNA-seq (10× Genomics’ 3’-capture method) compared to bulk RNA-seq with iPSCs, we were able to identify a putative “hot spot” where the expressed SNPs with allelic imbalance were clustered (70-80 Mb) on the X-chromosome in excitatory neurons across all donor lines (Figure 5E, Supplementary Figure 5B, Supplementary Table 6). Interestingly, this hot spot region also encompassed *XIST*. Looking at the same region in iPSC cells, we identified SNPs associated with 12 genes (*DLG3*, *SLC7A3*, *ZMYM3*, *PIN4*, *RPS4X*, *FTX*, *JPX*, *TTC3P1*, *NONO*, *TAF1*, *SEPHS1P4*, *RPS6P26*) (Figure 3G, Supplementary Figure 3D, Supplementary Figure 5C, Supplementary Table 6). This suggests that *XIST*-associated silencing may be confined to specific regions of the X-chromosome.

To systematically identify genes consistently influenced by *XIST* across different donor lines in iPSC-derived neurons, we mapped ASE SNPs (*XIST*^+^/*XIST*^-^ ASE < 0.75 or > 1.25) to their corresponding genes. Among X-linked genes, 13 genes for excitatory neurons and 2 genes (*FTX*, *NAP1L3*) for inhibitory neurons were identified in at least 2 cell lines of which 1 gene (*TTC3P1*) was consistently found across all cell lines in excitatory neurons (Figure 5F, Supplementary Table 5). *TTC3P1* was among 8 genes (*AL135749.6*, *HDAC8*, *MIR325HG*, *ATRX*, *NLGN3*, *FTX*, *JPX, TTC3P1*) that were present in the hot spot region in the X-chromosome of excitatory neurons. For autosomal genes, 313 genes for excitatory neurons and 63 genes for inhibitory neurons were present in at least 2 cell lines, where 25 genes for excitatory neurons and 3 genes (*BX470102.3*, *CRABP1*, *S100A6*) for inhibitory neurons were shared across all cell lines (Figure 5F). When we combine the neuronal types, 22 of 332 autosomal genes and 5 of 14 X-linked genes have been implicated in neurodevelopmental disorders (Supplementary Tables 4, 5). Furthermore, these genes were overrepresented in biological processes related to postsynaptic signaling, brain morphogenesis, and CNS neuronal development (Figure 5G). These observations raise the possibility that *XIST* erosion is associated with altered regulation of neurodevelopmental genes, although direct functional effects remain to be determined.

### XCI erosion in iPSC-derived neurons has an effect on common transcriptomic differential expression analysis

Given our observed effects of XCI erosion on ASE bias of both X-linked and autosomal genes in iPSC-derived neurons, we attempted to examine how variable *XIST* expression across samples may affect an empirical transcriptomic-wide differential expression (DE) analysis. Using the pseudobulk RNA expression values in our scRNA-data of iPSC-derived neurons from all four female-donor isogenic lines, we performed a linear regression analysis to identify *XIST*-associated gene expression, regressing out the effects of the LoF allele and cell line (see methods). We only identified 52 and 12 X-linked genes affected by *XIST* expression in excitatory and inhibitory neurons, respectively, where 9 genes were shared between the two neuronal subtypes (Figure 6A, 6B, Supplementary Table 7). The lack of *XIST*-associated autosomal genes was consistent with our above-described much weaker effect of XCI erosion on autosomal genes. Among the *XIST*-associated genes, we found a subset of genes implicated in neurodevelopmental disorders (NDD): 7 genes (*ALG13*, *CASK*, *GDI1*, *HNRNPH2*, *PAK3*, *EIF2S3, ZC4H2*) in excitatory neurons and 3 genes (*PAK3*, *DCX*, *ALG13*) in inhibitory neurons. Consistent with the gene set enrichment for iPSCs (Figure 3I), the enriched biological processes for these neuronal genes perturbed by *XIST* included TORC2 signaling, pyrimidine biosynthesis, heat acclimation, and regulation of acyl-CoA biosynthetic processes (Figure 6C).

**Figure 6:**
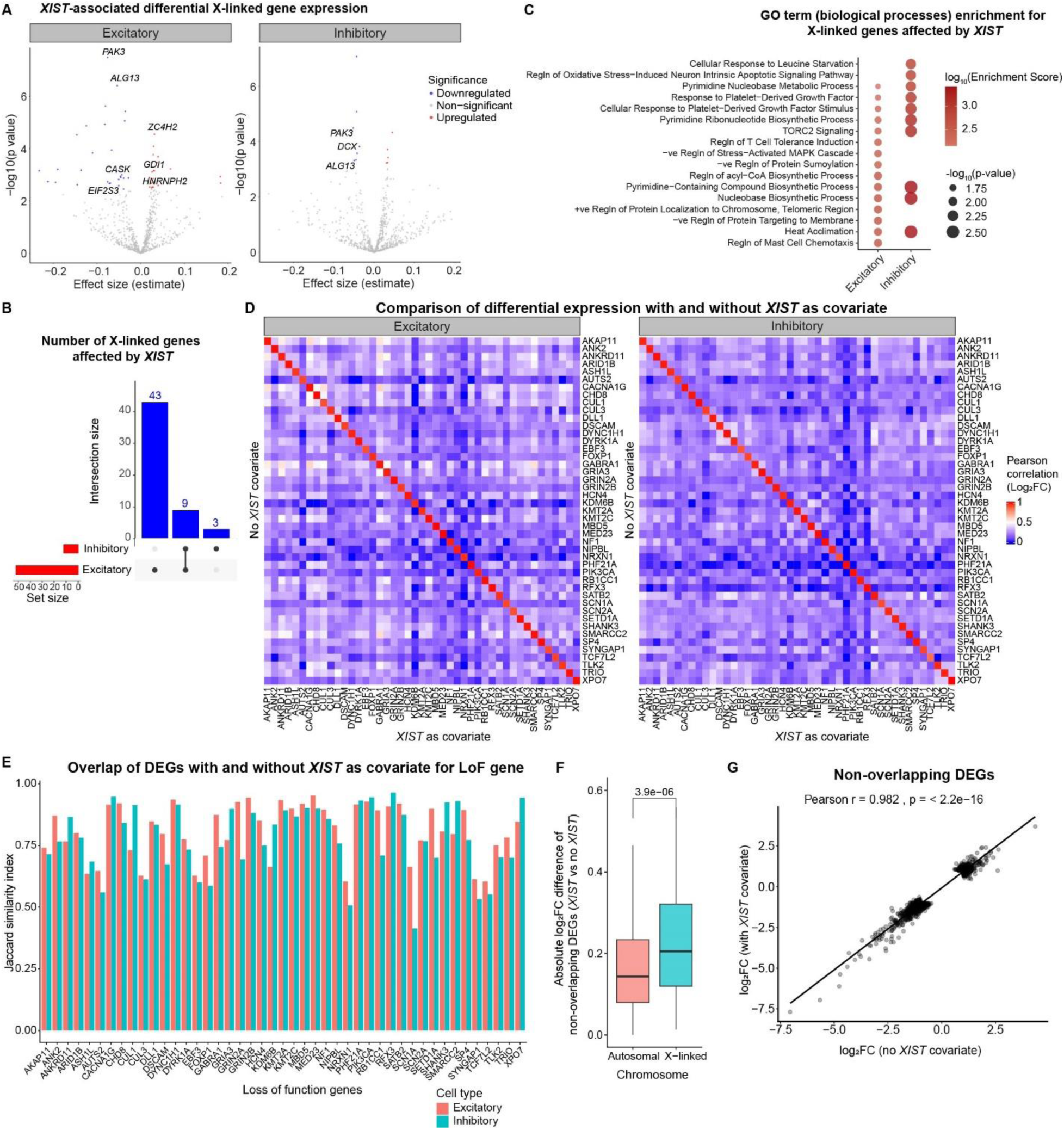
Impact of neuronal *XIST* expression on transcriptome-wide differential expression analysis. (A) Volcano plot showing X-linked genes with expression impacted by *XIST* expression based on standard linear regression. Genes with adjusted *p*-value < 0.05 are highlighted in red (upregulated) or blue (downregulated), of which NDD genes are labeled. (B) Upset plot showing *XIST*-associated genes from (A) in both cell types. (C) GO enrichment (biological process) for X-linked genes affected by *XIST* expression. (D) Pearson correlation of log_2_FC of gene expression, calculated using DESEq2 differential expression analysis for each iSTOP edited gene with and without *XIST* expression as a covariate. (E) Jaccard similarity index calculated using intersection of differentially expressed genes (*p*-value < 0.05 and log_2_FC < or > 1) identified using DESeq2 associated with each iSTOP edited gene with and without *XIST* expression as a covariate. (F) Comparison of autosomal and X-linked iSTOP edited genes based on the absolute log_2_FC differences (with *XIST* vs without *XIST* expression as a covariate) of non-overlapping DEGs (*p*-value < 0.05 and log_2_FC < -1 or > 1) with Jaccard similarity index < 0.8. Box represents IQR and whisker represents 1.5 * IQR. (G) Scatter plot showing correlation between log_2_FC (with *XIST* vs without *XIST* expression as a covariate) for non-overlapping DEGs (*p*-value < 0.05 and log_2_FC < -1 or > 1) with Jaccard similarity index < 0.8.

We next examined to what extent *XIST* erosion may confound any differentially expressed genes (DEGs) associated with LoF alleles. Using DESeq2^33^ to perform pseudobulk DEG analysis between neurons of isogenic lines for each LoF mutation with and without including *XIST* expression as a covariate. Because of the small number of DEGs that have an adjusted *p* value < 0.05, for comparison purpose we used a relaxed DEG cut-off (nominal *p* value < 0.05 and FC > 2). We found that the log_2_FC of the DEGs for each LoF line, with or without *XIST* expression as a covariate, highly correlated (Pearson’s R > 0.85) (Figure 6D). The list of DEGs with or without *XIST* expression as a covariate was also mostly overlapping; however, there were still about 20% of LoF genes showing a lower Jaccard similarity index between 0.5 to 0.75 (Figure 6E). As expected from our observed stronger effect of *XIST* on X-linked genes, the non-overlapping X-linked DEGs (with *XIST* vs. without *XIST* as a covariate) exhibited a larger FC difference compared to those non-overlapping autosomal DEGs (Figure 6F, Supplementary Table 8). Notably, despite a sizable number of non-overlapping DEGs, their log_2_FCs across all assayed LoF genes with or without using *XIST* expression as a covariate were highly correlated (Pearson’s R = 0.98) (Figure 6G). Altogether, these results demonstrate a subtle but significant confounding effect of XCI erosion on common DEG analysis, highlighting the need of accounting for variable *XIST* expression across samples for data interpretation.

## DISCUSSION

*XIST* is a central regulator of X-chromosome dosage compensation in females, ensuring balanced gene expression between sexes. However, the effects of CRISPR editing on *XIST* expression in iPSCs and iPSC-differentiated neurons are not well characterized, limiting our understanding of X-linked regulation in these cellular models for brain disorders. Here, we systematically investigated a large number of CRISPR-edited iPSC lines of multiple donors and iPSC-derived neurons, providing an in-depth view of how gene editing and neuron differentiation influence XCI erosion in female cells. We also examined the *XIST*-induced ASE bias on specific X-linked and autosomal genes in both iPSCs and neurons, highlighting an epigenetic regulatory mechanism that may be conserved across different genetic backgrounds and during neuron differentiation from iPSCs.

Our study demonstrates that XCI erosion observed in iPSCs largely persists in differentiated neurons. Consistent with these findings, Raposo et al. (2025) reported that XCI erosion in hiPSCs is maintained following differentiation into multiple lineages, including trilineage commitment and cardiomyocytes^14^. However, we did observe a small number of instances in which isogenic lines that expressed *XIST* in the iPSC state lost expression in induced neurons, or conversely gained *XIST* expression following neuronal differentiation, suggesting a possible stochastic nature of XCI in iPSCs that may be influenced by CRISPR editing and neural differentiation. Nonetheless, the overall persistence of XCI erosion in CRISPR-edited iPSCs and iPSC-derived neurons has important implications for disease modeling using iPSCs, as abnormal *XIST* expression has been associated with multiple diseases including autoimmune disorders and cancer and may influence the expression of NDD genes^34–36^. Supporting this, Topa et al. (2024) specifically focused on NDD genes and identified 17 X-linked genes affected by *XIST* in iPSCs and 10 in iPSC-derived sensory neurons, of which three genes overlapped between the two contexts^13^. In our induced neurons, four of these X-linked NDD genes (*PAK3*, *DCX*, *EIF2S3, ALG13*) were affected, with *EIF2S3* and *DCX* showing specificity to excitatory and inhibitory neurons, respectively.

XCI erosion has been reported to occur spontaneously with increasing passage number^2,10,37^. Contrary to these studies, we did not observe XCI erosion due to increased passage number, likely due to the narrow range (19-26) of passage numbers in our study. Alternatively, as Briggs et al. (2015) reported, cells at late passage may be more resilient to *XIST* loss^10^. Thus, differences in *XIST* expression and ASE observed in our study are unlikely to be influenced by passage number, allowing us to focus on *XIST*-dependent transcriptional effects. As *XIST* coats the X-chromosome to enforce transcriptional repression, *XIST*^-^ females showed the expected elevation of X-linked expression in both iPSCs and neurons. Notably, this increase was more pronounced in iPSCs, suggesting X-linked gene activation is partially constrained in differentiated neurons. Our analyses of known XCI-escaping gene expression and ASE bias in neurons also supported a stronger effect of XCI erosion in iPSCs than in neurons. However, despite weaker effects of XCI erosion in neurons, we observed a localized “hot spot” (70–80 Mb) on the X chromosome in excitatory neurons and iPSCs that exhibited allelic imbalance (Figure 3G, 5E, Supplementary Figures 3D, 5B). This region encompassed the non-coding genes *FTX* and *JPX* ^35,38,39^, which are known regulators of *XIST* activation, and overlapped the X-chromosome inactivation center region spanning Xq13-q21. Together, these findings support a model in which *XIST* loss leads to modest, gene-specific relaxation of X-linked repression in neurons.

Compared to the known effect of XCI erosion on X-linked genes in female iPSCs, its influence on autosomal genes has not been well established, especially in iPSC-derived neurons. Previous studies have shown *XIST* in female hESCs and hiPSCs spreads its repressive effect to some autosomal regions^12,13^. On the contrary, we did not observe significant repressive effects of *XIST* on autosomal gene expression by comparing bulk RNA levels between *XIST*^+^ and *XIST*^-^ iPSC lines or neurons. However, we did find modest negative expression correlation of *XIST* and autosomal genes in single neurons expressing variable levels of *XIST*. Effects of XCI erosion were also observed for autosomal genes in both iPSC and neurons from our ASE bias analysis by comparing *XIST*^+^ and *XIST*^-^ samples. Despite a much weaker effect of *XIST* expression on autosomal genes than X-linked genes, our DEG analysis using commonly used pipelines in iPSC-derived neurons supported a potential confounding effect of *XIST* expression on autosomal genes, highlighting the need to consider female *XIST* expression as a covariate in DEG analysis.

Our study has some limitations. Although we had large cohort of CRISPR-edited iPSC lines and their matching neuronal samples, only 4 female donor lines were included. Additional female donor lines would allow us to gain a more comprehensive view of the genetic loci exhibiting *XIST*-induced allelic imbalance of expression of both X-linked and autosomal genes. Moreover, our iPSC lines had a relatively narrow range of late cell passages (19-26); including unedited NTC lines of early passages and assaying more time points of neural differentiation would help reveal more of the dynamics of XCI erosion and its stage-specific effects on X-linked and autosomal genes. Nevertheless, our study represents the first extensive evaluation of XCI dynamics and its transcriptional impacts on X-linked and autosomal genes in the same set of CRISPR-edited iPSC lines and neurons. The observed persistent XCI erosion in iPSC-derived neurons and its confounding effect on commonly used DEG analyses have significant implications in improving the design and interpretation of hiPSC-based disease modeling of neurodevelopmental and other brain disorders.

## ACKNOWLEDGMENTS

This work was supported by RM1MH133065 grant, awarded to Z.P.P., J.D., and J.G.M. as a part of the SSPsyGene Consortium. We acknowledge the SSPsyGene Consortium members for the selection of NPD genes and supplying the hiPSC lines CW20107 (from CIRM) and KOLF2.2J (from the Jackson Lab). We further extend our gratitude to Molecular Genetics of Schizophrenia (MGS) investigators who contributed to the collection of samples that were used to derive MGS hiPSC lines. This study was further funded by NIH grants R01AA023797 and R01MH125528 (to Z.P.P.); R01MH106575, R01MH116281, and R01AG063175 (to J.D.); and RM1MH133065 (to Z.P.P., J.D., and J.G.M.).

## AUTHOR CONTRIBUTIONS

C.T. and E.K.O. performed the main analyses, created the figures, tables and wrote the manuscript. J.L. performed the main analyses. A.M., A.B., L.M., and D.S. performed the CRISPR-editing and hiPSC derivation and culture. S.D.d.L.G. and P.G. performed the neuronal differentiation. D.S. characterized and imaged the neurons. H.Z. prepared the library for scRNA-seq. S.Z., R.M.P., R.P.H., C.N.P., A.K., J.G.M., and A.R.S. assisted with project design and result interpretation. J.D. conceived and supervised the project and wrote the manuscript. All authors contributed to the manuscript writing and editing.

## RESOURCE AVAILABILITY

### Lead contact

Further information and requests for resources and reagents should be directed to and will be fulfilled by the lead contact, Jubao Duan.

### Materials availability

The hiPSC lines will be made available as part of the SSPsyGene Consortium to fulfill the NIMH (National Institute of Mental Health) material/data-sharing commitment.

### Data and code availability

All the reported data and code used during analysis, including hiPSC eSNP-Karyotyping results are deposited at https://doi.org/10.5281/zenodo.18776733. The RNA-seq data’s GEO accession numbers are GSE262442 and GSE322198.

## LIST OF SUPPLEMENTARY MATERIALS

Methods

Supplementary Text

Figures: S1 to S6

Tables S1 to S8

References (1-28)

## METHODS

### Source hiPSC lines

The Institutional Review Board (IRB) of Endeavor Health approved the study, confirming that the research met established ethical guidelines. As part of the SSPsyGene consortium agreement, CW20107 (from California Institute for Regenerative Medicine, CIRM) and KOLF2.2J (from Jackson Laboratory #JIPSC003902); an updated version of KOLF2.1J) were made available. The MorPhic consortium (morphic.bio) generated KOLF2.2J by precisely correcting heterozygous point mutation in COL3A1^1^. The other cell lines (CD14, CD19, GHI26, GHI49); were derived by the Duan lab^2–4^. CD14 and CD27 originated from lymphocytes collected from healthy participants enrolled in the Molecular Genetics of Schizophrenia (MGS) cohort^4^. A detailed list of the cell lines is provided in Supplementary Table 1.

### iSTOP gRNA design and cloning

The 66 genes reported here are part of the collection of an ∼250 NPD gene list curated as a part of the SSPsyGene Consortium (sspsygene.ucsc.edu). The optimal iSTOP-gRNA was designed and retrieved using iSTOP web-based tool (www.ciccialab-database.com/istop/#)^5^. Nonsense-mediated decay (NMD) capability was determined by verifying that the targeted nucleotide was positioned ≥55 nucleotides upstream of the final exon–exon junction^5,6^. gRNAs meeting thresholds of >50% predicted NMD efficiency and affecting >50% of annotated transcript isoforms were prioritized. When feasible, guide sites were chosen close to 5’ site to minimize effects of possible truncated protein. To minimize off-target editing, we selected only gRNAs without predicted off-target sites in the human genome, allowing up to two mismatches within the first eight bases^5,6^. For cloning the designed sgRNAs, pDT-sgRNA (Addgene #138271) vector was selected as a gRNA carrier. The vector was digested with BbsI-HF. After gel purification, a single strand oligo with a prefix, gRNA of interest, and a postfix was introduced into the vector backbone through Gibson assembly using NEBuilder® HiFi DNA assembly reagent. After cloning, miniprep plasmids were sequenced using M3 Rev primer. After genotyping, correct clones were expanded, and transfection-grade plasmid was prepared using an endo-free plasmid kit (QIAGEN:12362).

### hiPSC culture and transfection

The hiPSCs were maintained in mTeSRPlus (StemCell #100-0276) with Primocin (Invitrogen #ant-pm-1) on tissue culture plates coated with Matrigel (Fisher Scientific #08-774-552). Culture media was changed every other day and the colonies were passaged every 4-6 days when they reached confluency of 70%. For DNA transfections, cells were seeded at a density of 1.2-1.5×10⁵ cells per well in a 24-well plate (ThermoFisher #142475) using mTeSR Plus supplemented with 5 µM ROCK inhibitor (Tocris #1254). The following day, the medium was switched to antibiotic-free mTeSR Plus with 5 µM ROCK inhibitor after confirming adequate cell density (30–60% confluence) and viability. Each well was transfected with 750 ng of pEF-AncBE4max (Addgene #138270), 300 ng of pEF-BFP, and 300 ng of pDT-sgRNA carrying the variant-specific guide RNA using LipofectamineSTEM at a 1:2.5 DNA-to-reagent ratio. A total of 23 distinct pDT-sgRNAs were tested (one per well), with the one negative control in each batch. Culture medium was refreshed with standard mTeSR Plus containing Primocin at 24 hours and again at 48 hours post-transfection. 72 hours post- transfection, cells were prepared for single-cell sorting.

### Single hiPSC sorting and clonal culture

Singularized hiPSCs were sorted 72 hours post-transfection as individual cells into 96-well plates using a BD FACSAria Fusion, in mTeSR Plus supplemented with CEPT (1:10,000 chroman 1, emricasan, trans-ISRIB, and 1:1,000 polyamine supplement)^7^. Cells were dissociated with Accutase (StemCell #07920) for 7 minutes at 37°C, quenched with 1 ml mTeSR Plus with CEPT in 15 ml tubes, and centrifuged at 300 × g for 3 minutes. The resulting cell pellets were resuspended in 700μl mTeSR Plus with CEPT, filtered twice using 5ml Corning round-bottom tubes with 35 μm Nylon Mesh Screen filter caps (Falcon: #352235), and immediately processed for sorting. BFP⁺/GFP⁺ cells were sorted one per well for each experimental condition, while BFP⁺/GFP⁻ cells were sorted for the negative control. After sorting, cells were left undisturbed for 24 hours. Timed mTeSR Plus media additions were performed as follows: 50 µl at 48 hours and 72 hours post-sorting, 120 µl at 96 hours, and a refresh of 100 µl per well at 144 hours after aspirating 120 µl of existing media. Afterwards, 100 µl media was refreshed every other day for 10-14 days until colonies reached a suitable size for Sanger sequencing genotyping. All media handling was performed using an Integra’s MINI 6 electronic pipette.

### PCR and Sanger sequencing for LoF genotype confirmation

Once colonies reached an appropriate size, 8–12 colonies were selected per editing condition. Genomic DNA was extracted using QuickExtract DNA Extraction Solution (Fisher Scientific #NC9904870) and amplified by polymerase chain reaction (PCR) with locus specific primers. Sanger sequencing was performed on a 3730xl DNA Analyzer and the resulting data were analyzed using SeqScape v2.5 for automated genotype calling. Up to 4 colonies with confirmed homozygous editing or heterozygous editing (if no homozygous colonies were available) were expanded for RNA isolation and cell cryopreservation.

### RNA isolation for RNA sequencing (RNA-seq)

Based on sequencing results, two clones were selected and expanded from a single well on a 6-well plate. Once cultures reached ∼70% confluency, each clone was further expanded into two wells: one for cryopreservation and one for RNA isolation. For RNA isolation, cells were lysed in 800μl of Buffer RLT Plus (QIAGEN #1053393) and lysates were stored at -80°C until later processed using QIAGEN RNeasy Plus Mini Kit (QIAGEN #74134) according to the manufacturer’s instructions. Purified RNAs were then sent to Novogene for RNA-seq.

### Bulk RNA-seq data processing

FASTQ files were processed using Trimmomatic^8^ in order to remove adapter sequences, trim low-quality bases from read ends, and filter out reads that are too short after trimming, thus improving the quality of data before downstream analysis. A palindrome clip threshold of 30, simple clip threshold of 7, minimum read length of 50, and sliding window of 3 with an average quality of 18 was used. Following that, the sequences were aligned to a human reference genome using STAR (Spliced Transcripts Alignment to a Reference)^9^, and the output was used to generate a count matrix showing the number of reads per gene. The following parameters were used to run STAR on 48 threads: --quantMode GeneCounts, --alignSoftClipAtReferenceEnds No, --outFilterScoreMinOverLread 0.30, --outFilterMatchNminOverLread 0.30.

The package edgeR^10^ was used to convert the STAR output count matrix into a counts-per-million (CPM) table through the functions DGEList() with remove.zeros = T and cpm(). Only the subset of data with CPM greater than 1 was carried forward.

After adding the chromosome numbers for each gene, the data set was divided into X-linked (excluding *XIST*), autosomal, and mitochondrial genes. For each group, mean of all the reads for a given gene were compiled. From this, the average *XIST* expression could be compared between samples, experiment batches, and cell lines by simply grouping the data by the relevant characteristics. For further analysis, the data was also grouped based on sex and *XIST* expression. An *XIST* mean reads value greater than 1 was considered *XIST*^+^ while less than or equal to 1 was considered XIST-.

### CellNet analysis for pluripotency

R package CellNet^11^ was used for pluripotency assessment in each edited iSTOP hiPSC line. Gene × sample count matrix generated using edgeR was loaded, and a normalized count matrix with log-transformed, library size adjusted CPM values was generated using calcNormFactors(), estimateDisp(), and cpm(). A random forest classifier was then constructed using the built-in CellNet model. Finally, the likelihood scores for each sample–cell type pair were calculated by passing the normalized matrix through the classifier, and the results were visualized as hierarchically clustered heatmaps.

### Immunofluorescence staining for hiPSCs

iPSCs were dissociated using Accutase (Innovative cell technologies #AT-104) and plated onto Matrigel-coated (Corning #354234) 12 mm 1.5 image-grade glass coverslips placed in a 4-well plate. Cells were maintained in mTeSR Plus medium (Stem cell technology #100-0275) until medium-sized colonies formed. Culture medium was aspirated and cells were briefly washed with 1× PBS before fixation with 4% PFA at room temperature (RT) for 15 minutes. Following fixation, coverslips were washed 3 times with 1× PBS. Samples were then incubated in blocking buffer containing 4% bovine serum albumin (BSA; Sigma #A3803), 0.1% Triton X-100 (Fisher BioReagents #BP151-500), in PBS for 1 hour at RT. Primary antibodies (1 μg/ml) were then incubated for 1.5 hours. Samples were washed three times in PBS (5 minutes each) and incubated with the respective secondary antibodies (1:1000) for 1 hour at RT protected from light. After three additional PBS washes (5 minutes each), samples were incubated with DAPI (0.5 μg/ml; Fisher Scientific #EN62248) in PBS for 10 minutes at RT. The samples were then washed one final time in PBS and mounted onto glass slides using diamond anti-fade solution. The following antibodies were used: rabbit anti-Oct4 (1:250) (abcam #ab181557), mouse anti-SSEA4 (1:250; abcam #ab16287), goat anti-Human Nanog (1:20; R&D Systems #AF1997), donkey anti-Rabbit IgG Alexa Fluor Plus 647 (Fisher Scientific #A32795), donkey anti-Mouse IgG Alexa Fluor Plus 594 (Fisher Scientific #A32744), and chicken anti-Goat IgG Alexa Fluor 488 (Fisher Scientific #A21467). Confocal images were acquired using a Nikon C2 Confocal Microscope with a 20× objective.

### Lentivirus generation

Lentiviral vectors were generated by transfecting HEK293T cells with 3rd generation lentivirus packaging plasmids (pMDLg/pRRE, VSV-G, and pRSV-REV) with the desired vectors using lipofectamine 3000 (Invitrogen #L3000015). Culture media was collected 48 hours after transfection and centrifuged at 300 × g to remove cellular debris. The clarified supernatant was concentrated using a lentivirus precipitation solution (Alstem #VC100) with overnight incubation at 4°C, according to the manufacturer’s instructions. Samples were centrifuged for 30 min at 1,500 × g, and the resulting viral pellet was resuspended in mTeSR^+^ media, aliquoted, and stored at −80°C until use.

The following plasmids were used: pMDLg/pRRE (Addgene #12251), pRSV-Rev (Addgene #12253), pCMV-VSV-G (Addgene #8454), FUW-M2rtTA (Addgene #20342), FUW-TetO-Ngn2-P2A-puromycin (Addgene #52047), FUW-TetO-Ascl1-T2A-puromycin (Addgene #97329), FUW-TetO-Dlx2-IRES-hygromycin (Addgene #97330), and FSW-TdTomato (Addgene #197033).

### Neuron differentiation and co-culture

To generate neurons, hiPSCs were dissociated using Accutase (Innovative Cell Technologies, #AT-104), counted, and plated at 250,000 cells per well in Matrigel-coated 6-well plates (Corning #354234) in mTeSR Plus medium (StemCell Technologies #100-0275) supplemented with CEPT cocktail (Chroman 1, 50 nM; MedChemExpress #HY-15392; Emricasan, 5 µM; SelleckChem #S7775; Polyamine supplement, 1×; Sigma-Aldrich #P8483; trans-ISRIB, 700 nM; R&D Systems, 5284).

Cells were transduced at the time of plating with the following lentiviral vectors: i) Ngn2 + rtTA was added for excitatory neuron differentiation^12^, ii) Ascl1 + Dlx2 + rtTA was added for inhibitory neuron differentiation^13^. On day 1 post-infection, the media was changed to Neurobasal (Gibco #21103-049) supplemented with B27 (Gibco #17504044) and glutaMAX (Gibco #35050061).

Doxycycline (2 µg/mL; MP Biomedicals #198955) was added from day 1 to day 7 to induce transcription factor expression. Selection was performed from day 2 to day 6 using puromycin (2 µg/mL; Sigma-Aldrich #P8833) for excitatory neurons, or puromycin (2 µg/mL) plus hygromycin (100 µg/mL; Sigma-Aldrich #H9773) for inhibitory neurons.

On day 6, primary mouse glia was plated onto Matrigel-coated plates; for high content imaging, we plated 10,000 cells per well for a 96-well plate, for scRNA-seq assays we plated 100,000 cells per well for a 12-well plate. On the same day, excitatory neurons were transduced with a lentiviral vector expressing tdTomato under the human synapsin (hSyn) promoter to enable discrimination from inhibitory neurons during imaging analyses.

On day 7, induced neurons were dissociated with Accutase, counted, and replated onto glia-coated wells in Neurobasal medium supplemented with 5% FBS (R&D Systems #S11550) and CEPT cocktail. For high-content imaging, 15,000 excitatory and 7,000 inhibitory neurons were seeded per well in 96-well plates. For scRNA-seq experiments, 700,000 excitatory and 300,000 inhibitory neurons were plated per well.

On day 8 media was changed to neurobasal (with B27 and GlutaMAX) 5% FBS with BDNF (10ng/ml, PeproTech #10781-164), GDNF (10ng/ml, PeproTech #10781-226) and NT3 (10ng/ml, PeproTech #10781-174); Cytosine β-D-arabinofuranoside (AraC 2-4μM Sigma-Aldrich #C1768) was added to the media to stop glia proliferation. Half the media was replaced every 5 days with fresh neurobasal 5% FBS supplemented with BDNF, GDNF, and NT3. On day 35, the high content imaging plates were washed 2 times with PBS 1× and fixed with 4% PFA for 30 minutes. Cells were left in PBS 0.02% sodium azide until staining.

For scRNA-seq experiments, cells were dissociated on day 30 using Accutase for 30 minutes to obtain a single-cell suspension. Cells were filtered with a Flowmi cell strainer (Bel-Art #H13680-0040), centrifuged at 300 × g, and resuspended in Fluent Cell Buffer (Fluent Biosciences #FB0002440) for cell counting and viability assessment. Cells were then resuspended at the appropriate concentration in Fluent Cell Buffer prior to fixation. Cells were fixed using a methanol–DSP fixation protocol (Thermo Fisher Scientific #A35393) following the manufacturer’s instructions. Briefly, cells were incubated with methanol-DSP for 30 minutes, the reaction was quenched with 1 M Tris (pH 7.5). Fixed cells were stored at -20°C before downstream processing.

### High-content imaging for iPSC-derived neurons

Fixed iPSC-derived neurons were permeabilized in 1× PBS containing 0.5% Triton X-100 for 15 minutes at RT and then blocked with 3% BSA with 0.1% Triton X-100 in 1× PBS for 1 hour at RT. Primary antibodies were applied for 1.5 hours at RT. Samples were then washed 3 times 0.1% PBST (1× PBS + 0.1% Triton X-100) for 5 minutes each, followed by incubation with secondary antibodies (1:1000) for 1 hour at RT protected from light. After two additional washes with 0.1% PBST, samples were incubated with DAPI (0.5 μg/ml, Fisher Scientific #EN62248) for 10 minutes at RT. Plates were stored in 0.05% sodium azide in PBS at 4°C and equilibrated to RT prior to imaging. The following antibodies were used: goat anti-tdTomato (1μg/ml)(Fisher Scientific, 50-167-1115), chicken anti-MAP2 (1:5000, Sigma Aldrich #AB55543), donkey anti-Goat IgG Alexa Flour 568 (Fisher Scientific #A11057), and donkey anti-Chicken IgY Alexa Flour 647 (Fisher Scientific #A78952).

Images were acquired using the ImageXpress Micro Confocal High-Content Imaging System (Molecular Devices, San Jose, CA) with a 20× objective. DAPI, Texas Red, and Cy5 channels were used for fluorescence acquisition. Each well of the 96-well plate was imaged at nine sites per well, with 10 z-stacks collected at 1 µm step size. The 20× objective has a pixel size of 0.6842 µm² and a 60 µm pinhole.

### Single cell RNA sequencing library construction

Single cell RNA seq libraries were prepared using Illumina Single Cell 3’ RNA Prep T2 kit following vendor’s instructions. Briefly, mature neuronal culture was treated with Accutase at 37°C for 30-45 minutes. Cells were filtered with a Flowmi cell strainer (Bel-Art #H13680-0040), centrifuged at 300 g, and resuspended in Fluent Cell Buffer (Fluent Biosciences #FB0002440) for cell counting and viability assessment. Cells were then resuspended at the appropriate concentration in Fluent Cell Buffer prior to fixation. Cells were fixed using a methanol–DSP fixation protocol (Thermo Fisher Scientific #A35393) following the manufacturer’s instructions. Briefly, cells were incubated with methanol–DSP for 30 min, the reaction was quenched with 1 M Tris (pH 7.5). Fixed cells were stored at −20 °C before downstream processing. Following fixation, plates were shipped overnight to the Endeavor Health Research Institute (Evanston, IL). Later, fixed cells were resuspended in resuspension buffer (3× SSC, 1muM DTT, 0.2U/ul Rnase inhibitor, 1% BSA) and counted. For each bead’s aliquot, we loaded 5,000 fixed cells to target 2,000 cells per capture in the final library. cDNA quality was examined through Agilent bioanalyzer before proceeding to library construction.

### Single cell RNA sequencing (scRNA-seq) processing

Raw sequencing libraries were aligned to a hybrid human–mouse reference (GEX GRCh38 and GRCm39) using PIPseeker v3.3.0^14^. The Sensitivity-4 count matrix generated by PIPseeker was imported into Seurat to create the initial object^15^. Species identity was determined by calculating, for each cell, the proportion of human and mouse transcripts. Cells with ≥85% human-mapping transcripts were classified as human and retained for downstream analysis; mouse cells and putative doublets were excluded. The human-cells enriched object was further filtered to remove low-quality droplets and potential multiplets using the following criteria: > 500 nCount_RNA < 40,000, > 1000 nFeature_RNA < 7,500 & percent.mt < 10. Gene expression values were log-normalized using a scale factor of 1,000. All genes were used for scaling and variable stabilization. Cells were then clustered using the Leiden algorithm^16^. All sequencing batches and libraries were integrated using Harmony^17^ to correct for batch effects while preserving biological variation. Cell clusters were annotated based on established marker gene expression (neurons: *hg-MAP2*, excitatory neurons: *hg-SLC17A6*, inhibitory neurons: *hg-GAD1* and *hg-GAD2*, astrocytes: *hg-GFAP*, and mouse astrocytes: *mm-Gfap*). Additionally, we performed differential marker analysis for each cluster. Clusters in which the majority of marker genes were of mouse origin were removed. Additionally, clusters lacking clear marker signatures, were excluded from further analysis. Chromosomal metadata were added to the Seurat object using gene annotations from the GENCODE Release 46 (GRCh38.p14) GTF file^18,19^. Based on this annotation, genes were categorized as autosomal (chromosomes 1–22) or X-linked (X-chromosome) for downstream analyses. Depending on the expression of “*hg-XIST*”, “*hg-RSP4Y1*”, and “*hg-USP9Y*” we confirmed the identity of male and female cell lines. A clear expression difference of “*hg-XIST*” allowed us to distinguish the female libraries into *XIST*^+^ and *XIST*^-^.

We generated pseudo-bulk count matrices for the integrated scRNA-seq data using the AggregatedExpression() function from Seurat, where the cells were grouped by cell identity, library and batch. To remove batch effects, we ran negative binomial regression using the ComBat-seq() function from the R package sva with batch as a factor^20,21^. The resulting combat adjusted pseudobulk matrices were normalized using the DGEList() and CalcNormFactors() functions from the edgeR^10^ package. Counts per million (CPM) tables were then calculated using the cpm() function. To remove lowly expressed genes, we computed the mean CPM across all samples and retained only genes with a mean CPM > 1. For this new pseudobulk matrix, female samples with *XIST* CPM > 20 were categorized as *XIST*^+^, while those with *XIST* CPM < 20 were categorized as *XIST*^-^.

### Linear modeling of *XIST* expression in relation to X-linked and autosomal gene expression

To assess how *XIST* influences the expression of autosomal and X-linked genes in the scRNA-seq data, we restricted the analysis to *XIST*^+^ libraries for each cell type and only selected cells with “*hg-XIST*” > 0. We performed linear regression for each cell using the lm() function, modeling *XIST* expression as a function of the mean expression of X-chromosome genes (excluding *XIST*) and autosomal genes. For the bulk RNA-seq data, log₂-transformed mean read counts were analyzed using lm() to relate XIST expression to the average expression of X-linked genes (excluding *XIST*) and autosomal genes.

### Gene-wise analysis of *XIST* effects in pseudobulk samples

For female pseudobulk samples, linear models were fitted for each sample using lm() in R, with cell line and LoF gene included as covariates. Gene-wise regression was performed across all genes to identify genes whose expression was associated with *XIST* levels.

### Pseudobulk RNA-seq differential gene expression analysis

We evaluated how adjusting for *XIST* expression influences differential gene expression results by comparing two DESeq2 models^22^ using the DESeqDATASetFromMatrix() function. The first model included LoF gene, cell line, and log-transformed *XIST* CPM as covariates, whereas the second model omitted *XIST* expression.

To reduce noise from lowly expressed genes, we filtered out genes with < 10 total counts across all samples. An initial DESeq2^22^ run (DESeq()) was performed to calculate summary statistics, including the base mean expression for each gene. The dataset was filtered to retain only genes with a base mean ≥ 5, and the differential expression analysis was re-run on this filtered dataset to obtain final results.

We also evaluated overlap in significantly differentially expressed genes by computing the Jaccard similarity index for genes with *p*-value < 0.05 and absolute log_2_FC > 0.1 across all LoF genes and cell types. For LoF genes with similarity index < 0.8, we identified non-matching genes which were defined as genes that were significant in only one of the two models. These non-matching genes were categorized as autosomal or X-linked, and their log₂FC values from the *XIST*-adjusted and unadjusted models were extracted.

### Comparison of the change in gene expression in *XIST*^+^ and *XIST*^-^ females to the males

Combat corrected pseudobulk and bulk RNA-seq samples were used to compare the expression of escape genes in *XIST*^+^ and *XIST*^-^ females across cell lines. After converting the filtered counts matrix with DESeqDataSetFromMatrix(), standard analysis with the DESeq() function was completed^22^. DGE analysis was performed to compare the foldchange of autosomal and X-linked genes separately in *XIST*^+^ and *XIST*^-^ females to the males with no additional covariates. The result from the above differential expression analyses were used to make the escape gene heatmap plots. List of escape genes were extracted from a published study^23^.

### Allele specific expression analyses

All individual cells with XIST expression > 0 in the integrated Seurat object were marked as XIST^+^ and XIST^-^ otherwise. After organizing cell barcodes by XIST expression status (XIST^+^ or XIST^-^), all BAM files were then subsetted with the filterbarcodes command from Sinto (version 0.9.0, timoast.github.io/sinto). The resulting BAM files for each group were then converted back to FASTQ format using Picard’s SamToFastq tool.

Fastq files generated using Picard^24^ (for scRNA-seq) and Trimmomatic (for bulkRNA-seq) were aligned to the human genome (hg38) using Burrows-Wheeler Aligner. Duplicate reads were removed using Picards’s MarkDuplicates, and final BAM files for each library were generated by adding read-groups using AddOrReplaceReadGroups.

For bulk RNA-seq, BAM files were grouped by cell line and *XIST* expression status (*XIST*^+^ or *XIST*^-^). For scRNA-seq, BAM files were also grouped by cell type, resulting in distinct cell line × cell type × *XIST* status groups. To ensure comparability, BAM files were selected such that each group contained an equal number of samples with similar read depth. The number of bulk RNA-seq samples per cell line were CW20107 (n=9), CD14 (n=12), GHI126 (n=12), and GHI49 (n= 4). For scRNA-seq, the sample numbers per cell type and cell line were as follows: for excitatory neurons: CW20107 (n=4), GHI126 (n=5), GHI49 (n= 3), and CD14 (n=8); for inhibitory neurons: CW20107 (n=4), GHI126 (n=5), GHI49 (n=3), and CD14 (n=6). These selected BAM files were then used to generate VCF files using bcftools mpileup.

For *XIST*^-^ samples, SelectVariants from GATK^25,26^ was used to subset the VCF files to include only the heterozygous biallelic SNPs using the following parameters: --restrict-alleles-to BIALLELIC, --exclude-non-variants, --exclude-filtered, -select-genotype isHet==1. Next, a read counts per allele for ASE analysis was performed using ASEReadCounter^27^ with –min-mapping-quality = 10 and --min-base-quality = 2. For each group, a combined ASE bed file was created. Using the genomic position of the SNPs, the generated count table was merged for *XIST^+^* and *XIST^-^* samples. For each unique genomic position, the reference (refCount), alternate (altCount), and total counts (totalCount) were summed across all the entries. In order to remove low coverage data, SNPs with 0 total counts were removed. Finally, depending on the autosomal or X-linked chromosome analysis, the contig was filtered based on autosomal or X-linked analysis. The ASE ratio was calculated by dividing the sum of refCount by sum of totalCount. Bed files of these SNPs per cell line (and per cell type for scRNA-seq) were then exported to be used with GATK’s CollectAllelicCounts to analyze *XIST*^+^ samples. For the *XIST*^-^ group, we applied a threshold based on group size, retaining SNPs with a total read count ≥ (number of libraries per group * 5).

To compare the distribution of ASE ratios between *XIST*^+^ and *XIST*^-^ groups within each group, we generated density plots. Only nonzero *XIST*^+^ SNPs that could also be found in the filtered *XIST*^-^ data set were used. The minor allele count for each SNP was defined as the minimum of total reference and alternate read counts for each SNP. The ASE ratio was then calculated by dividing the minor allele count by the total read count at that SNP.

SNPs were considered biallelic if they had a *p*-adjusted value greater than or equal to 0.05 and ASE otherwise. From the merged table for each cell line, the biallelic SNPs were extracted for both *XIST*^+^ and *XIST*- groups. Among ASE SNPs, those with an *XIST*^+^ to *XIST*^-^ ASE ratio < 0.75 or > 1.25 were considered to have allele expression influenced by *XIST*. Using the position information, GENCODE Release 46 annotation gtf was used to map the gene for each *XIST*^-^influenced ASE SNP. NDD genes were obtained from Geisinger database^28^ and paired with the mapped gene lists.

### Statistical Analyses

All statistical analyses were performed in R version 3.6.3. Pairwise comparisons were assessed using the Wilcoxon rank-sum test, and global differences across groups were evaluated using the Kruskal–Wallis test. Multiple-comparison correction for pairwise tests was applied using the Benjamini–Hochberg false discovery rate (FDR) procedure where applicable. *XIST* CPM values were adjusted for cell line using lm() in R and compared with passage number, pluripotency score, and neuronal differentiation score. For all other analyses, unadjusted CPM values were used. Correlation analyses were performed using Pearson’s correlation coefficient.

## SUPPLEMENTARY TEXT

Supplementary Table 1: Metadata for bulk and single cell RNA sequencing libraries.

Supplementary Table 2: iPSC cells autosomal gene list for ASE SNPs with *XIST*^+^/*XIST*^-^ ASE ratio either below 0.75 or above 1.25

Supplementary Table 3: iPSC cells X-linked gene list for ASE SNPs with *XIST*^+^/*XIST*^-^ ASE ratio either below 0.75 or above 1.25

Supplementary Table 4: Neuronal autosomal gene list for ASE SNPs with *XIST*^+^/*XIST*^-^ ASE ratio either below 0.75 or above 1.25 with NDD genes

Supplementary Table 5: Neuronal X-linked gene list for ASE SNPs with *XIST*^+^/*XIST*^-^ ASE ratio either below 0.75 or above 1.25 with NDD genes

Supplementary Table 6: Hot spot ASE SNPs with *XIST*^+^/*XIST*^-^ ASE ratio either below 0.75 or above 1.25 for excitatory neurons and iPSC cells

Supplementary Table 7: XIST-associated gene list determined by standard linear regression for iNs

Supplementary Table 8: Non-matching genes for LoF with Jaccard index less than 0.8 and their log_2_FC

**Supplementary Figure 1:**
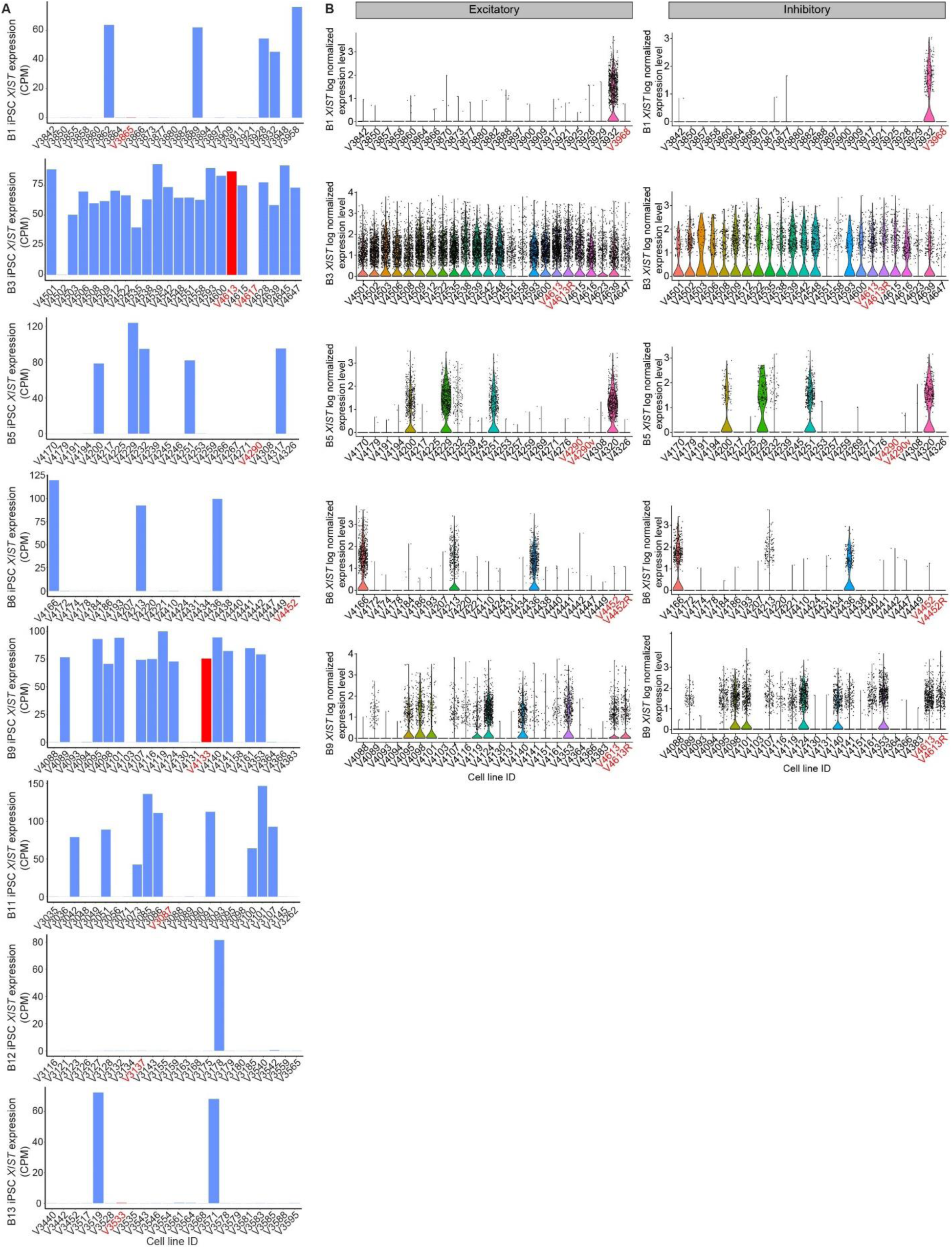
Expression of *XIST* across batches in bulk and single cell RNA sequencing. (A) *XIST* expression of each isogenic iPSC line for different editing batches in bulk RNA sequencing. CPM, counts per million reads. (B) Violin plot showing normalized expression of *XIST* in excitatory and inhibitory neurons of each isogenic line. Non-targeting control (NTCs, i.e., unedited lines) are labeled in red. Note that batch 1 NTC lines V3865 in (A) and V3968 in (B) are from 2 different iPSC clones, as well as batch 9 NTC lines V4133 in (A) and V4613 (and replicate V4613R).

**Supplementary Figure 2:**
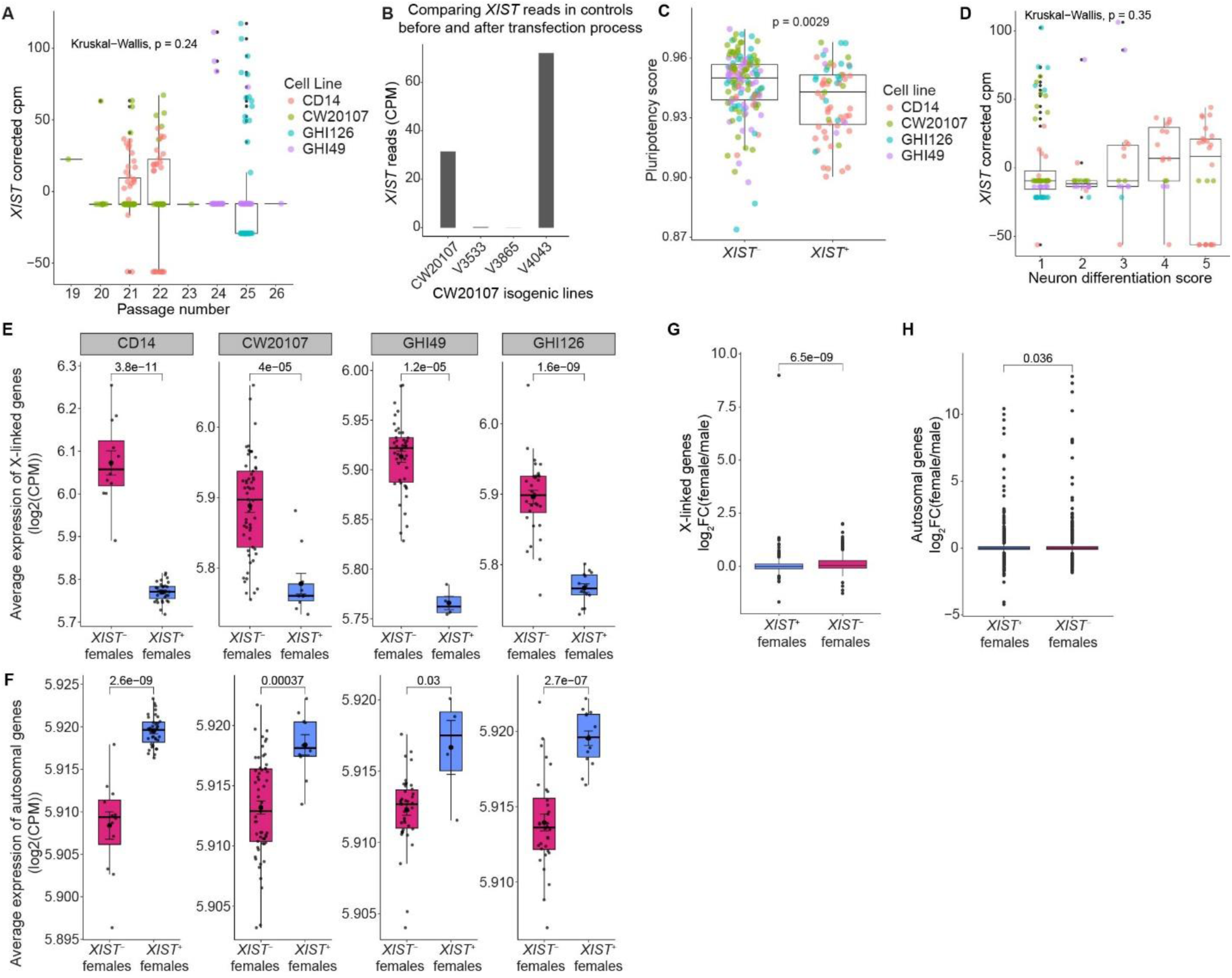
*XIST* expression across cell lines, passage number, editing status, differentiation score, and transcriptional profile. (A) Box and whisker plot comparing *XIST* reads adjusted for cell line using a linear model across passage number. (B) Bar plot showing expression of *XIST* (CPM, counts per million reads) in CW20107 line (original iPSC vial) and in its unedited NTC lines (V3533, V3865, and V4043). NTC lines have gone through the transfection and editing process without adding sgRNAs. (C) Comparison of pluripotency score between *XIST*^-^ and *XIST*^+^ female iPSC lines. Pluripotency score was calculated from RNA-seq data using CellNet. Wilcoxson *p*-value is shown. (D) Comparison of *XIST* reads adjusted for cell line using a linear model across iPSC lines with different category of neuron differentiation difficulty scores. (E-F) Box and whisker plot showing comparison of average expression levels of X-linked (E) and autosomal (F) genes for isogenic iPSC lines from four different female donor lines. (G-H) Box and whisker plot showing comparison of expression levels (normalized by male expression) of X-linked (G) and autosomal (H) genes for all isogenic iPSC lines. Box and whisker represent IQR and 1.5*IQR respectively. *XIST*^+^, with detectable *XIST* expression; *XIST*^-^, without detectable *XIST* expression.

**Supplementary Figure 3:**
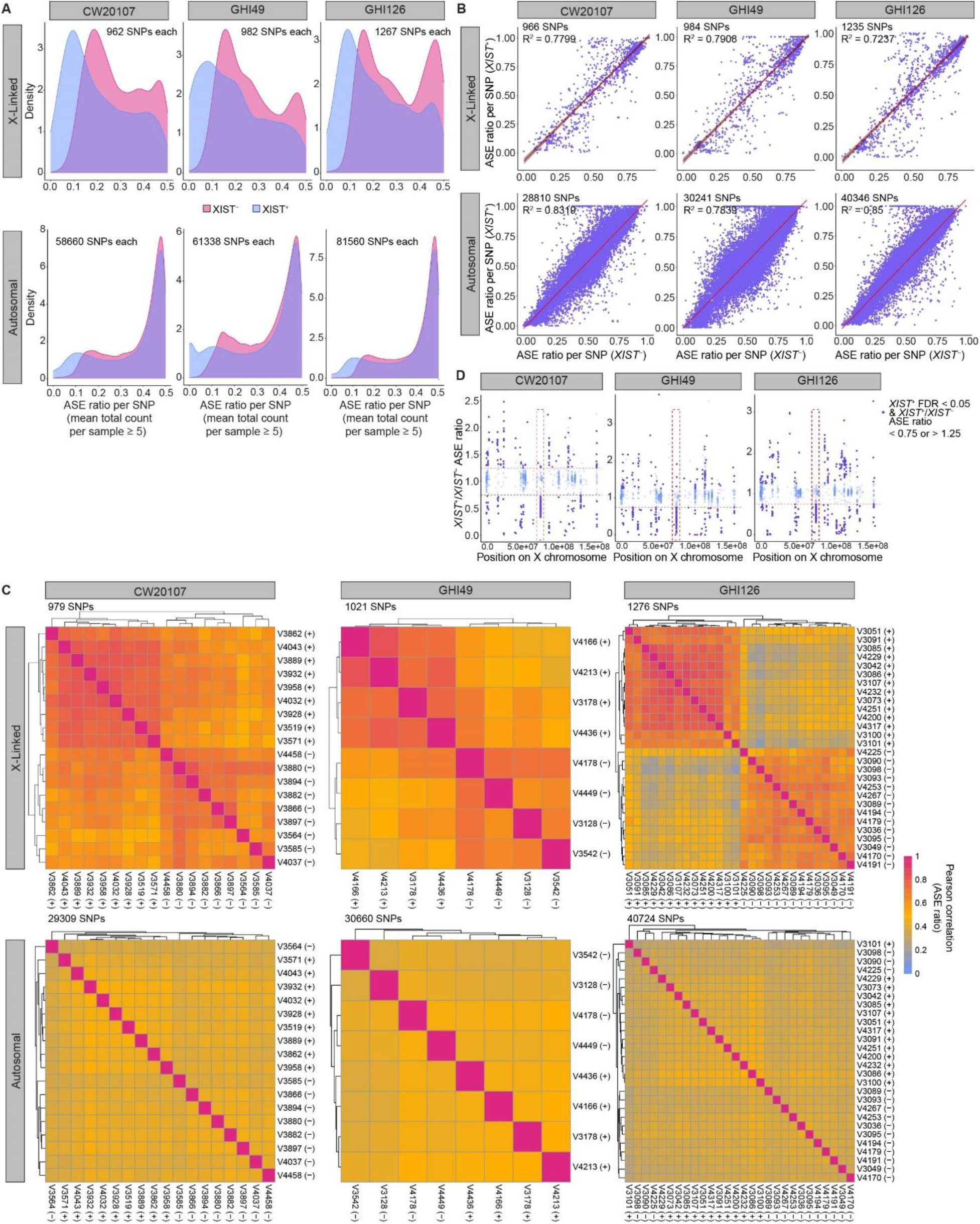
Comparison of SNP allelic imbalance of gene expression between different *XIST* status across iPSC cell lines. (A) Density plot showing the distribution of ASE ratio for X-linked (top) and autosomal (bottom) SNPs in the three female iPSC lines (not shown in main figure) with *XIST*^-^ or *XIST*^+^ status. (B) Scatter plot to compare the degree of allelic imbalance between *XIST*^+^ and *XIST*^-^ isogenic lines derived from the 3 female donors as in (A). Each dot represents one SNP. (C) Heatmap comparing Pearson’s correlation of SNP ASE ratio (ref Count / total Count) between isogenic iPSC lines with different *XIST* status (*XIST*^+^ or *XIST*^-^) for each donor line. (D) Positional scatter plot showing the distribution of *XIST*^+^/*XIST*^-^ ASE ratio for SNPs across the X-chromosome in different donor iPSC lines. Horizontal red dashed line highlights SNPs with *XIST*^+^/*XIST*^-^ ASE ratio < 0.75 or > 1.25 whereas vertical red dashed box highlights the 70-80Mb hot spot of allelic imbalance.

**Supplementary Figure 4:**
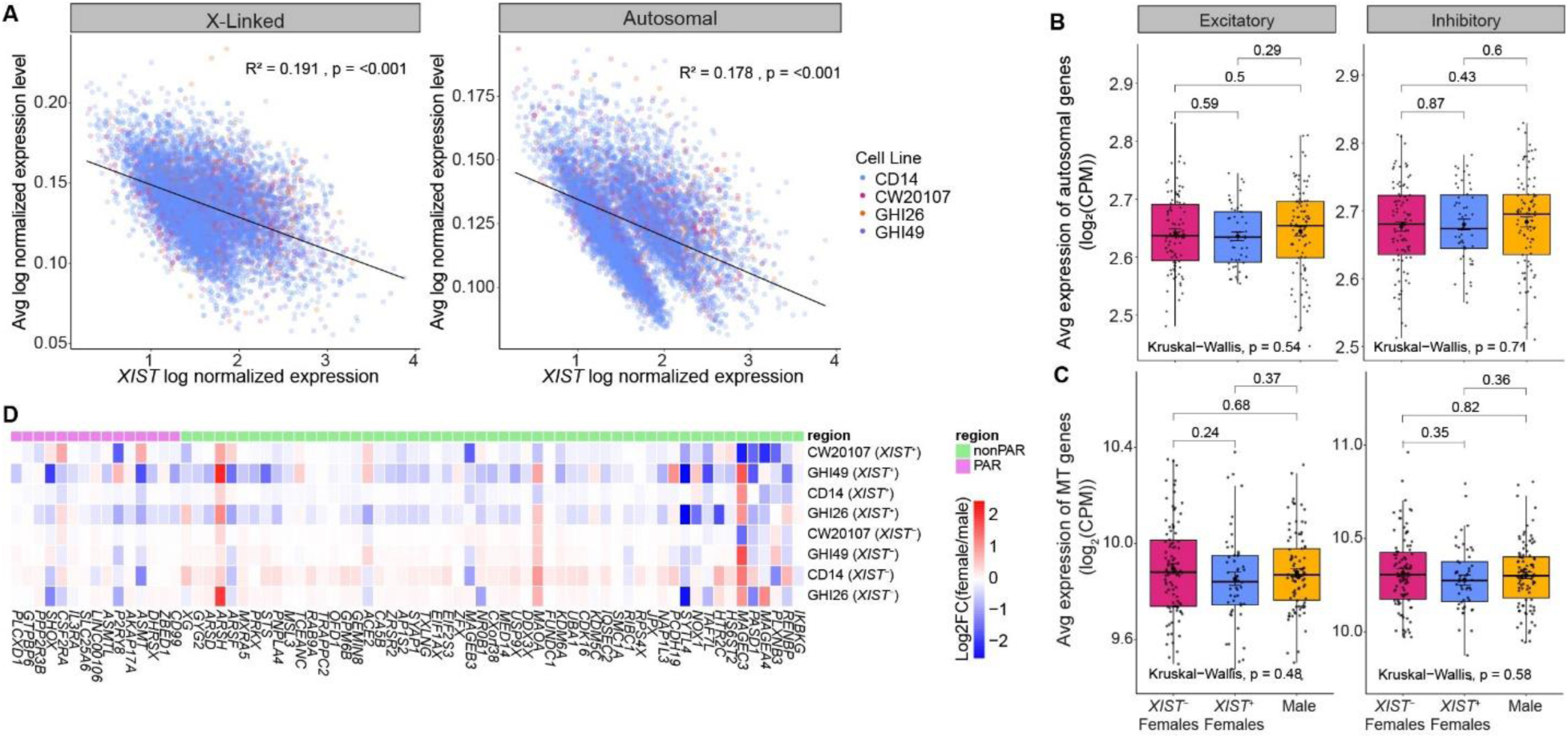
*XIST* expression impacts the expression of X-linked genes in iNs. (A) Linear regression plots showing the correlation of log-normalized *XIST* expression to average expression of X-linked genes and autosomal genes in inhibitory neurons. Each dot is a single cell where the colors denote different cell lines. (B-C) Box-and-whisker plots comparing the expression of (B) autosomal genes and (C) mitochondrial genome-encoded (MT) genes in isogenic iPSC lines between male and *XIST*^+^ and *XIST*^-^ females. Each dot represents pseudobulk data for an isogenic cell line. Box and whisker represent IQR and 1.5 * IQR respectively. (D) Heatmap showing the expression levels (normalized to male) of the reported XCI-escape genes (N = 70) in inhibitory neurons of female isogenic iPSC lines with different XCI status. PAR, pseudoautosomal regions of X-chromosome.

**Supplementary Figure 5:**
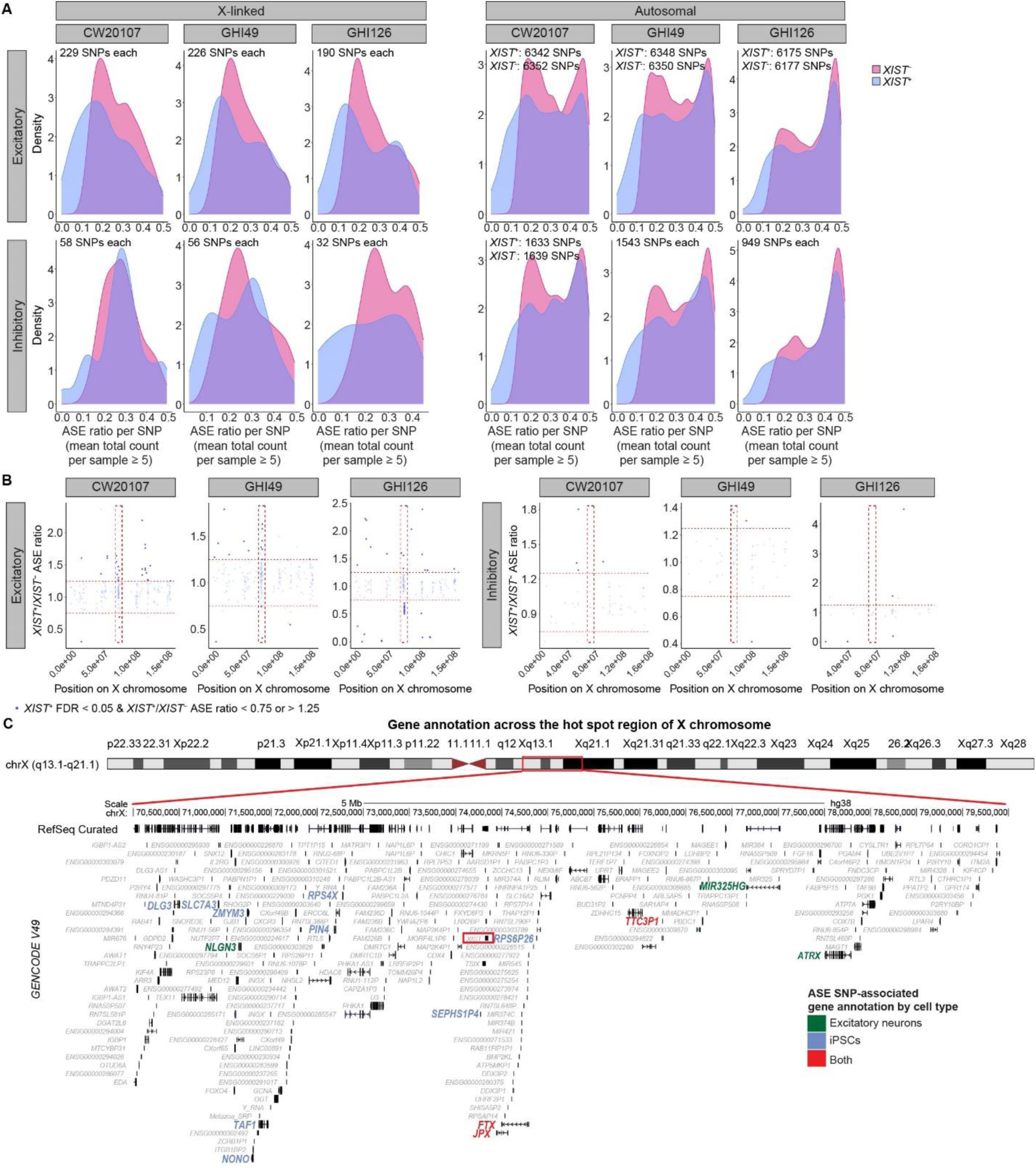
SNP allelic imbalance caused by XCI erosion in iPSC-derived neurons. (A) Density plot showing the distribution of ASE ratio for X-linked (left) and autosomal (right) SNPs between XIST- and XIST+ female iPSC-derived neurons for 3 different donor lines. (B) Positional scatter plot showing the distribution of XIST+/XIST- ASE ratio for SNPs across the X-chromosome of different cell lines and neuronal subtypes. Horizontal red dashed line highlights SNPs with XIST+/XIST- ASE ratio < 0.75 or > 1.25 whereas vertical red dashed box highlights the 70-80Mb hot spot of allelic imbalance. (C) Visualization of gene annotations across the 70–80 Mb region of the X-chromosome using the UCSC Genome Browser. The locus containing XIST is marked with a red box.

**Supplementary Figure 6:**
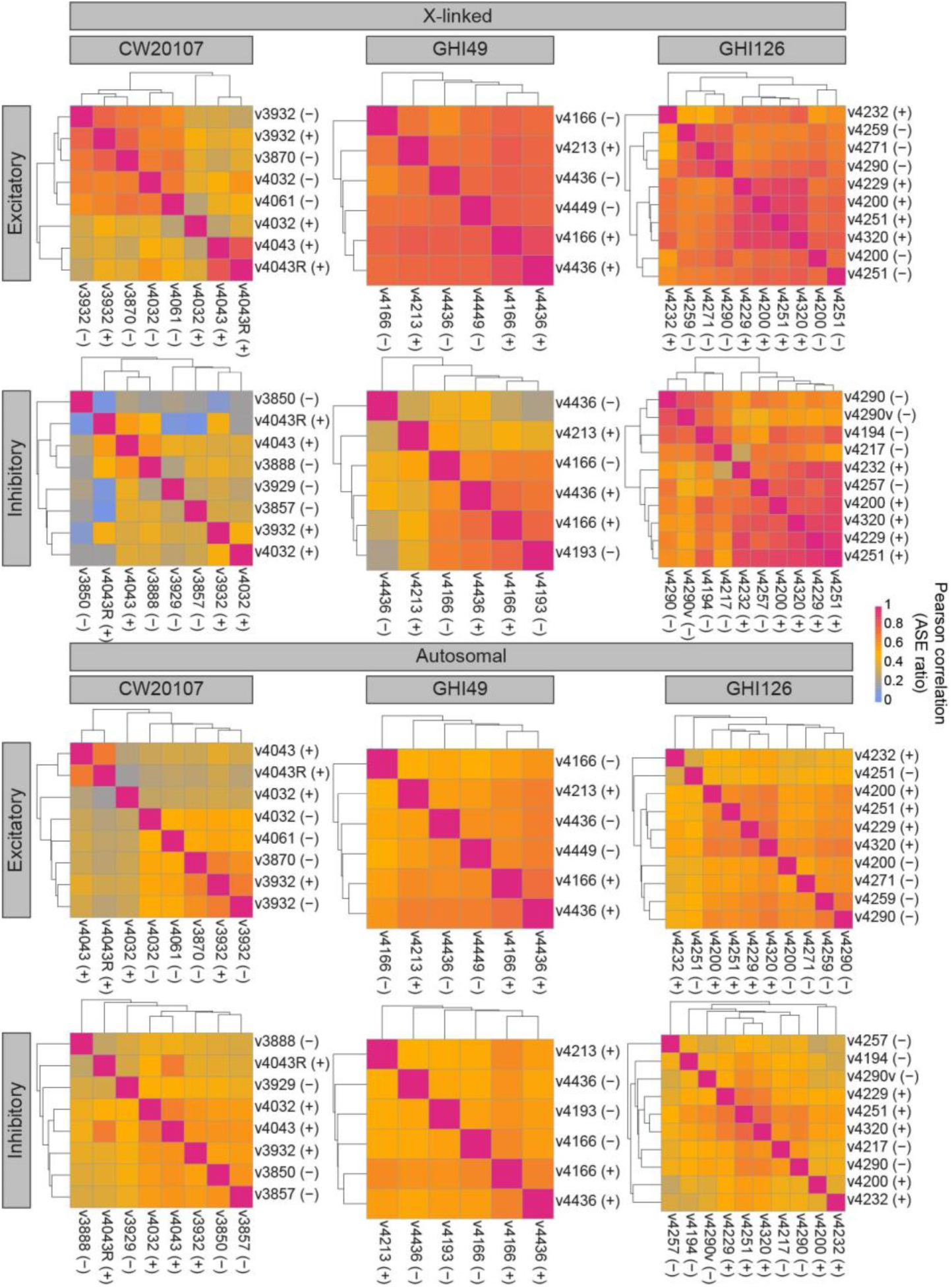
Pairwise Pearson’s correlation heatmap of SNP ASE ratio in iPSC-derived neurons between different lines. Correlation was calculated for X-linked and autosomal SNP ASE ratios (ref Count/total Count) in both excitatory and inhibitory neurons of isogenic lines with different XCI status (*XIST*^+^ or *XIST*^-^) for each donor line.

